# Dysregulation of neuroproteasomes by ApoE isoforms drives endogenous Tau aggregation

**DOI:** 10.1101/2022.11.29.518293

**Authors:** V Paradise, M Sabu, J Bafia, NA Sharif, C Nguyen, KD Konrad-Vicario, Mukim R Dhanraj, X Wang, BT Corjuc, J Fu, G Maldonado, J Ndubisi, M Strickland, H Figueroa, D Almeida, B Hyman, DM Holtzman, T Nuriel, KV Ramachandran

## Abstract

Neuroproteasomes are a subset of 20S proteasomes that are localized to the neuronal plasma membrane and degrade newly synthesized proteins. To date, the molecular composition of neuroproteasomes is undefined, and moreover, whether neuroproteasomes can influence protein aggregation with relevance to neurodegenerative disorders remains unexplored. Using a Cre-dependent conditional knock-in mouse line to endogenously tag the proteasome, we find that neuroproteasomes co-purify with ApoE, the most significant risk factor for late-onset Alzheimer’s Disease (AD). We discover that neuroproteasome membrane localization is differentially modulated by ApoE isoforms (E4<E3<E2) *in vitro*, *in vivo*, and in human postmortem samples. We synthesized selective, neuroproteasome-specific inhibitors and discovered that neuroproteasome inhibition induces aggregation of endogenous mouse and human Tau, without the need for seeding or pathogenic mutations. Using hApoE-KI/hTau-KI crosses, we find that ApoE isoforms differentially shift the aggregation threshold for Tau. Neuroproteasome inhibition *in vivo* is sufficient to induce sarkosyl-insoluble and Thioflavin-S positive endogenous Tau aggregates in only three days, which are completely abrogated by co-application of cycloheximide. Newly synthesized Tau levels increase threefold after neuroproteasome inhibition, leading us to posit that newly synthesized Tau is uniquely susceptible to aggregation due to neuroproteasome dysfunction. Overall, our data define neuroproteasomes as a pivotal proteostasis mechanism underlying the formation of endogenous Tau aggregates, which is directly regulated by the largest genetic risk factor for late-onset Alzheimer’s Disease.

## Introduction

Protein homeostasis mechanisms define the cellular capacity to handle protein synthesis, folding, and clearance while minimizing the accumulation of deleterious protein misfolding and aggregation events. While many factors contribute to protein homeostasis, protein degradation serves as a critical and non-redundant determinant of cellular proteostatic capacity. In canonical models of protein degradation, proteins are either degraded by the ubiquitin-proteasome system or the endo-lysosomal and autophagic systems(*1, 2*). In contrast to the established paradigms of protein degradation, we recently described the neuroproteasome, which is localized to the neuronal plasma membrane, exposed to the extracellular space, and degrades intracellular substrates across the membrane(*3*). We interpret this to mean that the neuroproteasome is functionally transmembrane(*3*). In addition, neuroproteasomes are composed of the 20S core proteasome and lack the canonical 19S cap and therefore, cannot recognize ubiquitylated substrates or unfold them for degradation. Instead, neuroproteasomes co- or peri-translationally degrade newly synthesized proteins without the need for ubiquitylation(*4*). While multiple reports have demonstrated the existence of neuroproteasomes in diverse systems(*5*), the molecular composition of neuroproteasomes, the regulation underlying neuroproteasome membrane localization, and the function of neuroproteasomes *in vivo* are poorly defined.

Failures in protein homeostasis are linked to accumulation of protein inclusions and aggregates. Indeed, Alzheimer’s Disease (AD) and many other neurodegenerative diseases are linked by the accumulation of protein aggregates(*6, 7*). Tau aggregates are among the most common intracellular aggregates in AD and are characteristic of more than 20 neurodegenerative diseases, termed Tauopathies(*8*). Multiple mutations that render Tau more prone to aggregation have been identified in patients(*9*) and these have been extensively modeled in transgenic lines and iPSC-derived neurons(*10-13*). While these studies have provided important insights into the mechanisms of Tau aggregate clearance, genetic variants in *MAPT* only make up less than 5% of the total population with AD and Tauopathies(*14-18*). Instead, endogenous Tau aggregates are found in the absence of overexpression or mutations in *MAPT* in the majority of patients with sporadic AD(*19, 20*). The protein quality control mechanisms in neurons that prevent endogenous Tau from aggregating remain elusive.

Moreover, aggregated Tau can induce the further aggregation of otherwise normal Tau, creating a feed-forward prion-like propagation referred to as Tau seeding(*12, 21, 22*). Studies delineating the principles of Tau seeding and propagation have provided new insights into the development of Tau pathology. Yet, the mechanisms underlying the initial aggregate formation in this pathological context are unclear. Therefore, identifying mechanisms by which endogenous Tau aggregates form could provide a unique therapeutic opportunity for intervention before Tau seeds can promote further templating and propagation.

While failures of the endolysosomal and autophagic systems can exacerbate Tau aggregation(*23-26*), whether and how proteasomal dysfunction affects Tau aggregation is contested. Multiple reports demonstrate that proteasomes can degrade Tau *in vitro* and in heterologous cells (*27-30*). However, follow up studies demonstrated that proteasome inhibition in neurons neither increases Tau levels nor does it induce endogenous Tau aggregation(*31-34*). In fact, somewhat counterintuitively, proteasomal inhibition leads to the reduction of soluble Tau due to the compensatory induction of autophagy in neurons(*31-34*). However, proteasomes can be involved in clearing already aggregated Tau but require the help of ATPases such as VCP to unfold the aggregates(*35*). The mechanisms by which protein degradation machineries determine both the formation and clearance of endogenous Tau aggregates are still debated. The discovery of the neuroproteasome provides a new mechanism to interrogate how protein degradation could contribute to the formation and clearance of protein aggregates in neurons.

While the precise mechanisms underlying Tau aggregation remain a subject of intense investigation, one major genetic factor that influences Tau aggregation and many other phenotypes in neurodegeneration is ApoE(*36-38*). The most significant risk factor for sporadic Alzheimer’s Disease is the ApoE4 isoform of the ApoE gene, while the ApoE3 isoform is neutral and the ApoE2 isoform is protective(*39, 40*). The mechanisms by which ApoE isoforms confer dramatic differences in the risk for AD remains unclear, but it is well established that ApoE4 is associated with increased Amyloid β and Tau pathology(*37, 41, 42*). More specifically, it remains mechanistically vague how ApoE isoforms influence Tau aggregation and neurodegeneration. Gaining the biochemical and cell biological insights connecting ApoE isoforms and Tau turnover could be invaluable to revealing the mechanisms underlying neurodegeneration.

Here, we discover an unexpected link between ApoE and endogenous Tau aggregation through the neuroproteasome. Using a transgenic mouse line to endogenously and conditionally tag the 20S proteasome coupled with quantitative proteomics, we found that neuroproteasomes co-purify with ApoE. Neuroproteasome localization at the plasma membrane is differentially regulated by ApoE isoforms, with ApoE4 reducing localization and ApoE2 increasing localization relative to ApoE3, *in vivo*, in primary neurons, and in humans. Using an unbiased quantitative proteomic screen, we identify neuronal responses to neuroproteasome inhibition in the soluble and insoluble proteome. Neuroproteasome dysfunction leads to deficits in proteostasis and induces aggregation of endogenous Tau in primary neurons and *in vivo*. We find that ApoE4 reduces the threshold at which Tau aggregates by ∼ 25-fold compared to ApoE3 and ∼200-fold compared to ApoE2, thereby dramatically elevating risk for Tau aggregation due to neuroproteasome dysfunction. Neuroproteasome-inhibition-induced endogenous Tau inclusions migrate as a high molecular weight species, which is typically reported only in the AD brain or with seeding-competent Tau(*43-46*). Using quantitative phosphoproteomic analysis supported by imaging in primary neurons and *in vivo*, we find that neuroproteasome inhibition induces the phosphorylation of Tau at sites consistent with pathological aggregated Tau in the AD brain. We suggest that neuroproteasomes serve as a pivotal proteostasis factor, mechanistically distinct from canonical degradation systems, directly linking ApoE to the formation of endogenous Tau aggregates.

## Results

### Endogenous affinity tags on the 20S proteasome reveal that neuroproteasomes co-purify with ApoE and Lrp1

We sought to purify the neuroproteasome directly out of the mouse brain. Our attempts to do this out of wild-type (WT) tissue proved challenging due to lengthy preparations which caused low yields. To circumvent this, we built a transgenic mouse line to endogenously tag the 20S proteasome for affinity purification. The proteasome is comprised of the 20S catalytic core particle which, as part of the ubiquitin-dependent degradation system, can be terminally associated with either one or two 19S regulatory particles, referred to as the 26S (20S+19S) or 30S (20S + two 19S) proteasome respectively. The 20S core is characterized by four axially stacked heptameric rings: the outer alpha subunit rings which gate substrate entry and the inner rings which contain the catalytic beta subunits. The 19S caps contain subunits to bind ubiquitylated substrates and ATPases to unfold said substrates into the 20S core. Therefore, modifying the 20S with affinity tags allows us to isolate any complex containing the core proteasome, such as free 20S, singly-capped 26S, doubly-capped 30S proteasomes, or the neuroproteasome. We modified the C-termini of nearly every core proteasome subunit with either 6X-His or Strep-II tags and screened out constructs based on low expression and interference with proteasome function **(Fig S1A)**. Of the remaining two constructs, we found that only the C-terminal modification of *Psma3* was compatible with proper proteasome assembly and function and did not adversely affect cell health **(Fig S1B)**. Based on these data, we generated a 20S-FLAG transgenic line by adding flanking loxP sites surrounding the final exon (Exon 11) of the *Psma3* gene followed by the same sequence with a linker containing a 3X-FLAG tag (**Fig 1A**). By crossing 20S-FLAG transgenic mice with a line expressing Cre recombinase driven by the pan-neuronal Actlb6 (BAF53b) promoter(*47*), we generated mice that expressed FLAG-tagged proteasomes selectively in neurons (**Fig 1B**). Brains from 20S-FLAG/BAF53b-Cre mice and Cre-only littermate controls were processed and FLAG-tagged cytosolic proteasomes were isolated on beads with a nanobody against the FLAG epitope. Simultaneously, 26S proteasomes were isolated using the GST-UBL affinity isolation method(*48*) and all samples were immunoblotted using antibodies raised against multiple 19S and 20S subunits. We find that the FLAG tag is incorporated into the proteasome and we fail to detect any FLAG in the Cre-only littermate controls (**Fig 1B**).

**Figure 1:**
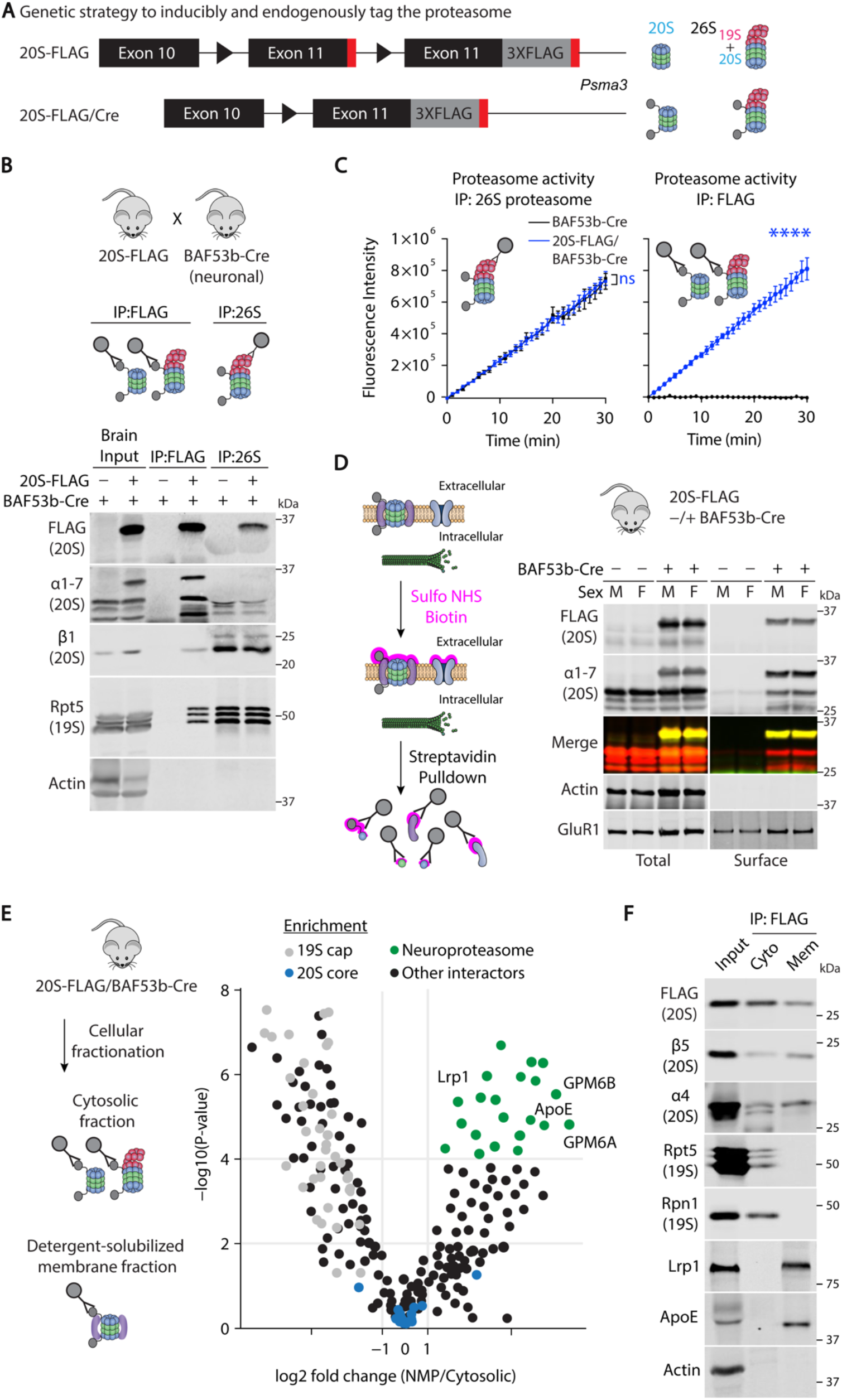
Neuroproteasomes co-purify with ApoE and Lrp1. **(A)** Schematic for endogenous and conditional tagging of the 20S proteasome with FLAG tag. (Top) Schematic of *Psma3* gene locus, endogenous Exon 11 containing the stop codon (red rectangle) was flanked with loxP sites (black triangles), and an insert containing 3x-FLAG tagged Exon 11 (gray) was inserted into the 3’ UTR of the gene. Transgene referred to as 20S-FLAG. (Bottom) Diagram of 20S-FLAG/Cre: in the presence of Cre, the native Exon 11 containing the stop codon was excised, driving the expression of FLAG tag from the endogenous *Psma3* locus. (Right, top) Schematics of proteasome complexes: 20S and 26S (20S + 19S regulatory particle). (Right, bottom) Schematics of proteasome complexes with 20S-FLAG tag (gray circles) after Cre recombination. **(B)** Proteasome purifications from the cytosol of brains from 20S-FLAG/BAF53b-Cre mice. (Top) 20S-FLAG mice crossed with pan-neuronal Cre driver line BAF53b-Cre to induce 20S-FLAG expression in neurons. (Middle) Diagram of proteasomal complexes isolated by different affinity methods; Immunoprecipitation (IP) using FLAG beads isolate FLAG-tagged 20S core particle as well as the 20S-containing 26S particle, whereas IP against the 19S only isolates 26S particles. (Bottom) FLAG and 26S IP from mouse brain cytosolic fractions immunoblotted using indicated antibodies. **(C)** Comparison of FLAG-20S proteasomes with unmodified native proteasomes. Catalytic activity of proteasomes isolated by 26S IP (left) and FLAG IP (right) from whole brain cytosol. Proteasomes were isolated as in (b) from BAF53b-Cre mice (black) or 20S-FLAG/BAF53b-Cre mice (blue). Catalytic activity of isolated proteasomes was measured by monitoring cleavage of model substrate, Suc-LLVY-AMC. Proteasome-dependent cleavage of the substrate increases AMC fluorescence. Data are mean ± SEM from three biological replicates. ****<0.0001 by Two-Way ANOVA. **(D)** Surface biotinylation to isolate surface-exposed proteins from 20S-FLAG/BAF53b-Cre hippocampi. (Left) Diagram of surface biotinylation and streptavidin pulldowns. Neuroproteasomes (20S surrounded by purple accessory proteins) and other membrane proteins (blue) depicted in membrane, cytosolic proteins such as microtubules (green) in intracellular space. Hippocampal tissues from 20S-FLAG mice alone or with BAF53b-Cre were dissected and subjected to surface biotinylation using amine-reactive sulfo-NHS-biotin (pink). Lysates incubated with streptavidin beads to pull down surface-exposed proteins. (Right) 20S proteasome signal in surface pulldown denotes the neuroproteasome. Streptavidin pulldown of surface proteins in 20S-FLAG male (M) or female (F) mice, -/+ BAF53b-Cre. Lysates (Total) and Streptavidin pulldowns (Surface) immunoblotted using indicated antibodies against the FLAG epitope tag, the 20S proteasome (α1-7), a cytosolic protein (Actin), and a membrane protein (GluR1). Merge indicates overlay (yellow) between FLAG (green) and α1-7 blots (red). **(E)** Co-IP and mass spectrometry (MS) analysis of 20S-FLAG proteasomes from cytosolic and membrane-enriched fractions. (Left) Cytosolic and membrane fractions were prepared from whole brains of 20S-FLAG/BAF53b-Cre and immunoprecipitates were isolated using anti-FLAG affinity beads. (Right) Samples were subjected to tandem mass tag (TMT) labeling and quantitative mass spectrometry. Differential levels of proteins enriched with membrane-associated neuroproteasomes vs. cytosolic proteasomes were analyzed and plotted as log2-fold change vs –log10(P-value). Green dots represent proteins enriched in the neuroproteasome IP, blue dots represent 20S core subunits, and gray dots represent proteins in 19S cap. **(F)** Validation of Co-IP proteomics data. 20S-FLAG Co-IP from cytosolic (cyto) and membrane (mem) fractions of 20S-FLAG/BAF53b-Cre mouse brain tissue (inputs). Fractions were immunoblotted using indicated antibodies.

We next measured the catalytic activity of the FLAG-tagged proteasome compared to unmodified proteasomes by monitoring the cleavage of a model substrate, Suc-LLVY-AMC(*49*). An increase in fluorescence reflects the proteasome-dependent cleavage of this LLVY substrate which mobilizes the free fluorescence of AMC. We find that the catalytic activity of isolated 26S proteasomes from mice expressing 20S-FLAG are indistinguishable from the unmodified proteasomes isolated from controls (**Fig 1C**). We find that FLAG affinity purified proteasomes are catalytically identical to 26S affinity purified proteasomes (**Fig 1C**). We fail to detect FLAG expression in the livers from 20S-FLAG/Baf53b-Cre mice, nor can we affinity isolate proteasomes using the FLAG epitope handle from the liver, indicative of the selective and inducible nature of the 20S-FLAG transgene (**Fig S1C, D**). Overall, we find that mice expressing 20S-FLAG appear indistinguishable from wild-type littermates. These extensive characterizations support the conclusion that epitope tagging of the 20S with FLAG is inert and does not disrupt native proteasome function.

To test whether the 20S-FLAG was efficiently incorporated into the neuroproteasome, we performed surface biotinylation experiments out of both primary neurons and hippocampal tissue from 20S-FLAG transgenic mice. This is a well-established method for measuring neuroproteasome surface localization(*3, 50, 51*). Hippocampi from 20S-FLAG/Baf53b-Cre mice were incubated with a cell-impermeable sulfonylated Biotin-NHS-Ester to label surface-exposed amines and surface proteins were pulled down on streptavidin beads. We observe strong FLAG expression in the surface fraction, indicating that the FLAG tag is incorporated into the neuroproteasome (**Fig 1D**). We fail to detect cytosolic proteins such as Actin in our surface fraction, validating our surface-labeling approach, and fail to observe FLAG expression in 20S-FLAG mice without Cre (**Fig 1D**). By blotting using an antibody which detects six of the seven alpha subunits of the 20S proteasome including α7, the subunit coded for by *Psma3*, we can clearly distinguish the modified α7 subunit overlaying with the FLAG signal, denoting modification of the endogenous protein (**Fig 1D**). Next, we used surface biotinylation to assess if our transgenic system was functional in primary cultures. Primary neurons from 20S-FLAG mice were cultured and transduced with AAVs to express Cre recombinase at days *in vitro* (DIV) two and then processed for surface biotinylation at DIV14 (**Fig S1E**). We did not detect FLAG without AAVs to express Cre, supporting our conclusion that the 20S-FLAG transgene is inducible and not leaky.

Next, we purified FLAG-neuroproteasomes to determine their composition. First, we prepared plasma membranes from 20S-FLAG/Baf53b-Cre brains and then gently extracted membrane proteins. We find our membrane preparations were enriched in membrane proteins such as GluR1 and depleted of cytosolic proteins like Actin (**Fig S1F**). We affinity isolated FLAG-neuroproteasomes from the membrane fraction as well as cytosolic FLAG-proteasomes and subjected both to quantitative tandem mass tag-based mass spectrometry analysis. As expected, we find common signatures of 20S subunits between both cytosolic proteasomes and neuroproteasomes and a significant depletion of 19S subunits in the neuroproteasome isolations. This is consistent with a lack of cytosolic subunits in our membrane preparations and validates our previous findings(*3-5*). We identified 64 proteins enriched in the neuroproteasome isolations which are depleted in the cytosolic fraction. This revealed significant enrichment of the GPM6 glycoprotein, which we previously described(*3*) (**Fig 1E**), and a surprising and strong enrichment of ApoE and the ApoE receptor Lrp1 (**Fig 1E, Table S1**). We do not observe other ApoE receptors, such as ApoER2, VLDLR, and LDLR in this analysis. We first validated these proteomic data by affinity isolating FLAG-neuroproteasomes as well as cytosolic FLAG-proteasomes and then immunoblotting these samples. Here, we validate our proteomic data by also identifying robust co-purification of ApoE and Lrp1 with the FLAG-neuroproteasome (**Fig 1F**).

### ApoE isoforms differentially modulate neuroproteasome surface localization *in vivo*

ApoE can be expressed as one of three isoforms in human: ApoE4, ApoE3, and ApoE2, each of which differs by only one or two residues. We next tested whether ApoE isoforms could differentially modulate neuroproteasome localization and function. We obtained human ApoE isoform knock-in (KI) mice, where the entire coding region for the *Apoe* gene was replaced with a cassette encoding the three human *Apoe* loci, generating a fully humanized ApoE protein(*52*). These mice therefore only express the ApoE2, ApoE3, or ApoE4 isoforms. hApoE-KI mice were perfused and then hippocampus and primary motor cortex (PMC) were microdissected and subjected to surface biotinylation and immunoblotting. We chose a limited survey of regions based on their differential susceptibility to accumulate pathological hallmarks of AD (hippocampus>PMC)(*53, 54*). We found a strong reduction of neuroproteasome surface localization in the hippocampus of ApoE4-KI mice, compared to ApoE3-KIs or ApoE2-KIs, measured by surface signal of 20S subunit β5 (**Fig 2A**). We observe a significant difference between ApoE3 male and female mice, where female ApoE3 mice have 50% fewer surface-localized neuroproteasomes than male ApoE3 mice in hippocampus (**Fig 2A**). We find this noteworthy in the context of the extensive literature on the influence of biological sex on AD pathogenesis(*36, 55-58*). When we monitor neuroproteasome localization in the PMC from the hApoE-KI mice, we find that neuroproteasome surface localization in ApoE4-KIs is reduced compared to ApoE2-KIs and we observe a trend, but no significant change in ApoE4-KI compared to ApoE3-KI (**Fig S2A**). These data may be attributable to differences in ApoE expression in different brain regions(*36, 52, 59*), a dimension which we have not systematically investigated.

**Figure 2:**
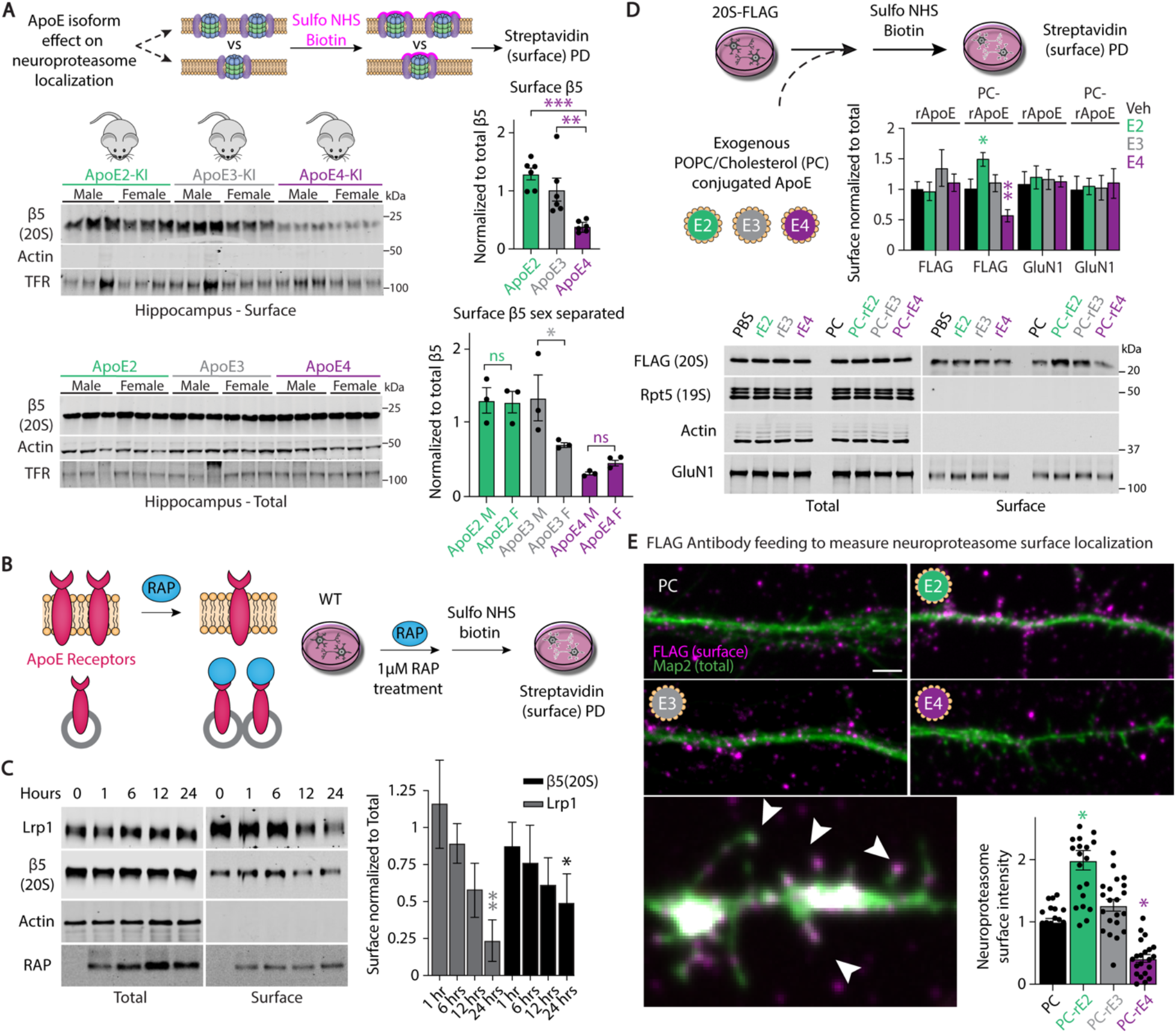
ApoE isoforms differentially modulate neuroproteasome localization in primary neurons and *in vivo*. **(A)** Surface biotinylation of hippocampi from hApoe-KI mice to assess neuroproteasome localization. Schematic (top) of surface biotinylation experiments from ApoE isoform knock-in (ApoE 2, 3, or 4-KI) mice to determine if neuroproteasome localization changes with ApoE genotype. Hippocampi (hippo) from ApoE2-KI (green), ApoE3-KI (gray), and ApoE4-KI (purple), male (M) and female (F) mice were subjected to surface biotinylation and streptavidin pulldown to isolate surface exposed proteins. Experimenters were blinded to genotype. Lysates (Total) and Streptavidin pulldowns (Surface) were immunoblotted using indicated antibodies. Quantification of surface β5 intensity was normalized to total β5 intensity. Data (right) are mean ± SEM normalized to ApoE3-KI, N=6 biological replicates per genotype (3 M, 3 F), ***p<0.001, **p<0.01 by One-Way ANOVA, *p<0.05 by Two-Way ANOVA Tukey’s Multiple Comparisons Test. **(B,C) (B)** Schematic of surface biotinylation of Receptor-Associated Protein (RAP)-treated neurons. Influence of RAP (blue) on surface ApoE receptors (light red) is depicted. Gray circles represent endosomes. **(C)** DIV13-14 primary cortical neurons obtained from WT mice were treated with 1 µM RAP for indicated time subjected to surface biotinylation. 20S proteasome signal in surface pulldown denotes the neuroproteasome. Lysates (Total) and Streptavidin pulldowns (Surface) were immunoblotted using indicated antibodies. Quantification of surface Lrp1 and β5 intensity was normalized to corresponding total signal. Data (right) are mean ± SEM normalized to 0 hr condition. N=3 biological replicates, **p<0.01, *p<0.05 by One-Way ANOVA Tukey’s Multiple Comparisons Test. **(D)** Surface biotinylation of 20S-FLAG neurons treated with exogenous ApoE isoforms (top left). DIV2 primary cortical neurons from 20S-FLAG mice were transduced with Cre AAVs to drive 20S-FLAG transgene expression. At DIV13, neurons were treated with exogenous unconjugated recombinant ApoE isoforms (400 nM; rE2: green; rE3: gray; rE4: purple), or POPC/Cholesterol (PC)-conjugated (yellow shell) recombinant ApoE isoforms (400nM; PC-rE2: green; PC-rE3: gray; PC-rE4: purple) for 24 hours and subjected to surface biotinylation. Lysates (Total) and Streptavidin pulldowns (Surface) were immunoblotted using indicated antibodies. Quantification of surface FLAG and GluN1 intensities were normalized to corresponding total intensities. Data (above) are mean ± SEM normalized to corresponding vehicle control. N=3 biological replicates, **p<0.01, *p<0.05 by One-Way ANOVA Tukey’s Multiple Comparisons Test. **(E)** Antibody feeding of 20S-FLAG neurons treated with exogenous ApoE isoforms. DIV2 primary cortical neurons from 20S-FLAG mice were transduced with Cre AAVs to drive 20S-FLAG transgene expression. At DIV13, neurons were treated with 400 nM PC-E2 (green), PC-E3 (gray), and PC-E4 (purple) for 24 hours. FLAG antibodies were fed on live neurons to label surface FLAG-neuroproteasomes (FLAG Surface). Subsequently, neurons were fixed and stained for MAP2 (MAP2 Total). Quantification of surface FLAG fluorescence intensity was normalized to total MAP2 fluorescence intensity. Data (below) are mean ± SEM normalized to PC alone. N=3 biological replicates, n=21 quantified regions, *p<0.05 by One-Way ANOVA Tukey’s Multiple Comparisons Test. Scale bar=2µm.

Given the region-specific effect of ApoE isoforms on neuroproteasome expression *in vivo*, we approached the effect of ApoE isoforms and ApoE receptors in a more defined and tractable primary neuronal culture system. Lrp1 was the only ApoE receptor that we identified in our neuroproteasome IP/MS dataset and we also found ApoE isoform-dependent modulation of Lrp1 levels (**Fig S2B**). Instead of testing the role for Lrp1 specifically, we tested the role for ApoE receptors in modulating neuroproteasome localization using the well-established pan-ApoE receptor antagonist Receptor-associated protein (RAP) (*60-62*). We chose this strategy because Lrp1 knockouts still contain small amounts of Lrp1 remaining in sporadic sets of neurons and because of detrimental effects on neuronal health following Lrp1 knockdown(*63, 64*). We used a variant of RAP (stable RAP) which contains a stabilizing mutation to protect against pH-induced denaturation and maintains a higher percentage of internalized Lrp1, and presumably other ApoE receptors as well(*60*) (**Fig 2B**). We incubated primary neurons with purified stable RAP and measured surface protein levels using surface biotinylation and immunoblotting. We find that stable RAP rapidly decreases the surface localization of Lrp1, as expected (**Fig 2C**). We also find that stable RAP reduces surface localization of neuroproteasomes (**Fig 2C**). These data served as the first demonstration that ApoE receptors, such as Lrp1, can regulate neuroproteasome localization.

ApoE is a lipoprotein, and in the brain is thought to be released by glia in the lipid-bound form(*36, 52, 65, 66*). We therefore next leveraged the primary culture system to determine whether extracellular ApoE could modulate neuroproteasome localization and if the lipidation of ApoE was important for this modulation. We generated primary neurons from either 20S-FLAG transgenic animals or WT animals and conducted imaging and biochemical experiments to measure neuroproteasome localization. We conjugated recombinant ApoE isoforms with a mixture of POPC and Cholesterol (PC) and purified ApoE lipoproteins, referred to as PC-ApoE2, E3, and E4. These lipoproteins were validated by SEC and negative stain EM (**Fig S2C**) and have been demonstrated to resemble the endogenous ApoE lipoproteins observed in the brain(*67*). DIV13 primary neurons from 20S-FLAG transgenic mice were incubated with unconjugated or PC-conjugated ApoE for 24 hours and then subjected to surface biotinylation. We find that PC-ApoE, and not the unconjugated versions, modify neuroproteasome surface localization with the rank order PC-ApoE4<Vehicle,PC-ApoE3<PC-ApoE2 (**Fig 2D**). We find no change in other membrane proteins such as GluN1, which has previously been shown not to change in an ApoE isoform-dependent manner(*68, 69*). The lipoprotein-dependent effect on neuroproteasome localization requires ApoE, as the addition of just the liposomes without ApoE does not modify neuroproteasome localization (**Fig 2D**). We find similar results in primary WT neurons, suggesting that the observed ApoE-isoform dependent effect on localization is not an artifact of the FLAG epitope tag on neuroproteasomes (**Fig S2D**).

FLAG-neuroproteasomes are a useful tool to measure the surface localization of neuroproteasomes and we next took advantage of the epitope tag to test whether ApoE isoforms modified the subcellular distribution and localization of neuroproteasomes in intact neurons. We used antibody feeding onto live neuronal cultures from the 20S-FLAG mice transduced with AAVs to express Cre(*70, 71*). No staining was observed using secondary alone controls (**Fig S2E**) or when feeding an antibody against intracellular protein MAP2 (**Fig S2F**). After live cell anti-FLAG antibody feeding, neurons were fixed, permeabilized, and stained for total intracellular MAP2 to label dendrites. Consistent with this approach being a robust measure of neuroproteasome localization (**Fig S2G**), we observed punctal neuroproteasome localization when feeding anti-FLAG antibodies, of which more than 90% was within 2 μm of the MAP2 signal. We observe significant overlap of surface FLAG signal with MAP2-positive protrusions that resemble dendritic spines (**Fig 2E**). We observe that PC-ApoE2 increases neuroproteasome surface localization by nearly twofold and that PC-ApoE4 reduces neuroproteasome surface localization by nearly threefold compared to PC vehicle or PC-ApoE3 controls (**Fig 2E**). In total, these data recapitulate the observations from the hApoE-KI mice and support our hypothesis that extracellular ApoE isoforms can influence neuroproteasome localization and function.

### Neuroproteasome localization is reduced in vulnerable brain regions in the human brain and is further reduced by ApoE4 genotype

We next tested whether ApoE-dependent mislocalization of neuroproteasomes is observable in postmortem patient tissues. For these analyses, we studied a cohort of 19 individuals (14 ApoE3/3 and 5 ApoE4/4) from the Massachusetts General Hospital Brain Bank. The ApoE3/3 subgroup was subdivided into 5 negative diagnoses and 9 positive diagnoses for AD. Sex was evenly distributed in samples except for the ApoE4 cases which were all male. This design allows us to compare the effect of AD on neuroproteasome localization as well as the effect of ApoE genotype on neuroproteasome localization. We obtained material from the Brodmann Area (BA4 – primary motor area) and BA7 (parietal associative area). We analyzed these regions as a means of untangling how neuroproteasome localization changes in a limited survey of severely pathologically affected (BA7) versus less affected (BA4) regions of the brain. Tissue from BA7 contained severe tangle and amyloid pathology as well as neuronal loss whereas tissue from BA4 had moderate amyloid pathology but not tangle accumulation or neuronal loss. For simplicity, we will refer to BA7 as tissue with severe pathology and BA4 as tissue with mild pathology. We fractionated these samples to remove nuclei, mitochondria, and cytosol, enriching for plasma membranes and detergent-insoluble proteins. We monitored the success of our fractionation using LLVY-based proteasome activity assays out of the supernatant of our washes and continued washing our plasma membrane preparations until cytosolic proteasomes were undetectable (**Fig 3A**).

**Figure 3:**
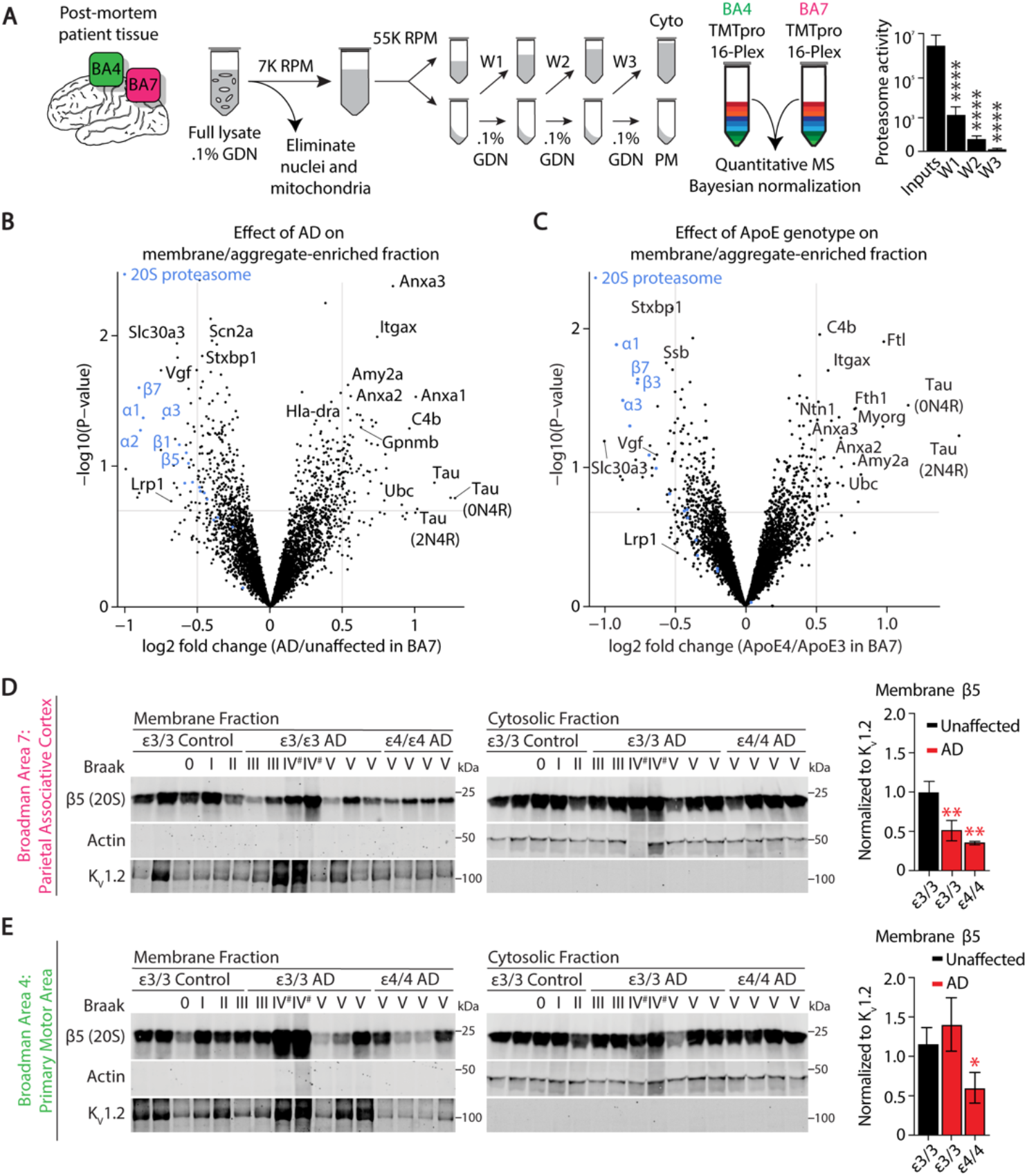
Neuroproteasome localization is reduced in AD-vulnerable brain regions in the human brain and is further reduced by ApoE4 genotype. **(A)** Schematic depicting steps to enrich plasma membranes from post-mortem human brain tissues. Post-mortem sections from Brodmann Area 7 (BA7, pink) and BA4 (green) were fractionated into crude membrane fractions, which were then washed 3 times (W1, W2, W3). Supernatants from washes were pooled to generate cytosolic fractions (Cyto). Membrane-enriched fractions also contain detergent insoluble protein (membrane/aggregate-enriched, PM). PM samples were labeled with Tandem Mass Tags (TMTpro) for quantitative MS. Cytosolic contamination assessed by the proteasome catalytic activity of the washes, graphed (right). N=32, ****p<0.0001 by Paired T-Test. Experimenters were blinded to sample identity. **(B)** Determining effect of AD on the plasma membrane and detergent-insoluble proteome. Differential enrichment of plasma membrane and detergent-insoluble proteins from BA7 of AD vs. unaffected patients was analyzed and plotted as log2-fold change vs –log10(P-value). Blue dots indicate 20S proteasome subunits, analysis was also done blinded to sample identity. **(C)** Determining effect of ApoE genotype on the plasma membrane and detergent-insoluble proteome. Differential enrichment of plasma membrane and detergent-insoluble proteins from BA7 in Apoε4/4 vs Apoε3/3 patients was analyzed and plotted as log2-fold change vs –log10(P-value). Blue dots indicate 20S proteasome subunits, analysis was also done blinded to sample identity. **(D)** Determining effect of AD and ApoE genotype on neuroproteasome membrane localization in BA7 (pink). Membrane and cytosolic fractions from BA7 from Apoε3/3 patients with AD (AD) and without AD (unaffected) and Apoε4/4 patients with AD were immunoblotted using indicated antibodies. Braak stages indicated above. (Right) Quantification of membrane β5 intensities were quantified and normalized to membrane loading control K_v_1.2. Data (right) are mean ± SEM normalized to ε3/3 unaffected. # indicates samples excluded from analysis on basis of high abnormal actin and K_v_1.2 signal. **p<0.01 by Two-Way ANOVA Fisher’s LSD Test relative to BA7 ε3/3 unaffected. **(E)** Determining effect of AD and ApoE genotype on neuroproteasome membrane localization in BA4 (green). Experimental setup is the same as described in (d), except BA4 was evaluated rather than BA7. (Right) Quantification of membrane β5 intensities were quantified and normalized to membrane loading control K_v_1.2. # indicates samples excluded from analysis on basis of high abnormal actin and K_v_1.2 signal. Data (right) are mean ± SEM normalized to ε3/3 unaffected. *p<0.05 by Two-Way ANOVA Fisher’s LSD test relative to BA4 ε3/3 unaffected.

We then subjected plasma membrane/aggregate fractions to quantitative TMT proteomics analysis. TMT proteomics enables unbiased quantification of the proteome and the ability to make quantitative comparison across samples. We compensated for brain region and individual variability in two ways: by first normalizing protein loading into the mass spectrometer and then normalizing the collected data against the total TMT signal per patient sample. This total TMT signal normalization provides the highest confidence metric for protein abundance normalization across all samples. Given the number of samples, we combined three TMTpro 16-plex experiments with a shared internal standard which enabled quantitation across TMT experiments. We first performed a series of quality control checks to ensure that these data provided reliable and reproducible measures. First, these data validate our plasma membrane enrichment protocol – we find a significant enrichment for membrane-bound and membrane-associated proteins and a strong depletion of cytosolic proteins, including the 19S cap (**Table S2**). Second, consistent with previous global proteomic analysis from AD brains, we find a significant increase of C4b(*72*), AnxA1(*73*), Gpnmb(*74*), and Tau (Mapt)(*75*) in AD brains compared to unaffected controls (**Fig 3B**). Third, we also find a significant decrease in Vgf(*76*) and Slc30A3(*77*) consistent with previous AD global proteomic datasets (**Fig 3B**). We expect to find aggregated proteins using this protocol since our plasma membrane enrichment method would also isolate large aggregates such as aggregated Tau in the AD brain.

After validating our approach, we next analyzed the effect of AD on neuroproteasome localization, measured by the relative quantity of 20S proteasome subunits in our quantitative mass spectrometry dataset. First, in comparing AD samples to cognitively normal controls, we find a striking reduction in nearly every 20S proteasome subunit in the plasma membrane fraction from BA7 tissue (**Fig 3B, Table S2**). We next sought to determine whether ApoE genotype influenced neuroproteasome localization in human tissue. Given the significant effect of AD on neuroproteasome localization, we constrained the analysis only to the effect of ApoE genotype on neuroproteasome localization by comparing the ApoE4/4 AD cohort against the ApoE3/3 AD cohort. We observe a clear trend of reduced 20S proteasome subunits in the membrane fraction in the ApoE4 genotype compared to ApoE3 genotype in the BA7 vulnerable brain region (**Fig 3C, Table S2**).

To validate our proteomics data, we monitored surface localization of neuroproteasomes by immunoblot analysis. We controlled for sample loading by normalizing against Kv1.2, a protein whose expression has been reproducibly demonstrated as unchanged in the AD brain(*78-80*). While this strategy is required for loading controls by immunoblot, these loading controls line up with the normalization strategy for mass spectrometry based on the total amount of TMT signal from each sample. First, we validate our proteomics data and find a twofold reduction of neuroproteasome surface expression in AD patients compared to controls in the BA7 region (**Fig 3D**). Moreover, we find a statistically significant reduction in neuroproteasome localization from ApoE4/4 tissue compared to ApoE3/3 unaffected tissue (**Fig 3D**). In contrast, we do not observe changes in total expression of the 20S proteasome from the cytosolic fraction, suggesting that the ApoE-isoform dependent changes we observe are specific to neuroproteasome localization and not a change in cytosolic proteasome levels. We find changes in Lrp1 concordant with that observed with neuroproteasomes (**Fig S3A**). However, given the strong effect of AD pathology on the global proteome (as well as on neuroproteasome localization), we next used the BA4 tissue to understand how ApoE isoforms could influence neuroproteasome localization with mild AD pathology.

In BA4 tissue, unlike BA7, we find comparable surface localization of neuroproteasomes between patients with and without AD (**Fig 3E, Fig S3B**). However, once stratified by ApoE genotype, we find that ApoE4/4 patients have a nearly 50% reduction in surface localization of neuroproteasome compared with ApoE3/3 patients (**Fig 3E**). Both of these conclusions are supported by analysis of proteasome subunits from unbiased proteomics datasets from BA4 tissue (**Table S2**). Consistent with these observations, using LLVY degradation experiments to measure the catalytic activity of the proteasome, we found a significant reduction in neuroproteasome activity in AD samples compared to unaffected controls and no change in cytosolic proteasome activity (**Fig S3C**). This demonstrates that ApoE4 can influence neuroproteasome localization in areas with mild AD pathology. Taken together, we make two conclusions based on these data: 1) that neuroproteasome localization is reduced in brain regions which have increased susceptibility to AD pathology and 2) that ApoE4 reduces neuroproteasome localization compared to ApoE3.

### Quantitative proteomics reveals that selective inhibition of neuroproteasomes induces accumulation of sarkosyl-insoluble Tau

What is the consequence of ApoE4-dependent reduction in neuroproteasome function, or more broadly, any reduction in neuroproteasome function? The most straightforward way to address this question is to leverage neuroproteasome-specific inhibitors. We previously accomplished neuroproteasome-specific inhibition using Biotin-Epoxomicin: an analog of the covalent and highly selective proteasome inhibitor Epoxomicin with an N-terminal Biotin moiety. This small modification renders Biotin-Epoxomicin cell-impermeable for up to an hour and does not change the selectivity or potency of Epoxomicin (*4*) (*81-84*). While time-limited, this approach found that generally, neuroproteasome substrates are co- or peri-translationally degraded and that flux through the neuroproteasome depends on neuronal activity(*3, 4*). However, the previous version of Biotin-Epoxomicin was not a usable tool to assess broad physiological functions of neuroproteasomes because the timescale was far too short and could not be used *in vivo*.

Based on principles of drug design, we considered that introducing a longer PEG linker between the N-terminus of Epoxomicin and biotin would decrease the permeability of biotin-epoxomicin and allow us to reveal long-term consequences of neuroproteasome inhibition(*85, 86*). We chose Epoxomicin because it leverages the unique chemistry of the catalytic threonines of the proteasome – the α-amino group of the threonine opens the epoxide of Epoxomicin which forms the final covalent morpholino adduct. The catalytic residues of Serine and Cysteine proteases do not have α-amino groups, which is the principle that renders Epoxomicin completely specific both against its target and relative to other proteasome inhibitors. These advantages explain why dozens of analogs of Epoxomicin have been made to the N-terminus without altering the specificity of the drug against the proteasome: no off-target effects of Epoxomicin or its analogs have been found in the two decades since its discovery (*83, 84, 87-89*). We synthesized click-able analogs of Epoxomicin (Azide-Epoxomicin) that would allow us to append a broad range of linkers (**Fig 4A**). We then conjugated Biotin-PEG24-Alkyne to Epoxomicin, producing Biotin-PEG24-Epoxomicin, or iBEp (**Fig 4A**). We observe efficient conjugation by HPLC analysis (**Fig S4A, B, C, D**). All molecules were HPLC-purified as single peaks and confirmed by LC/MS analysis (**Fig S4D, E**).

**Figure 4:**
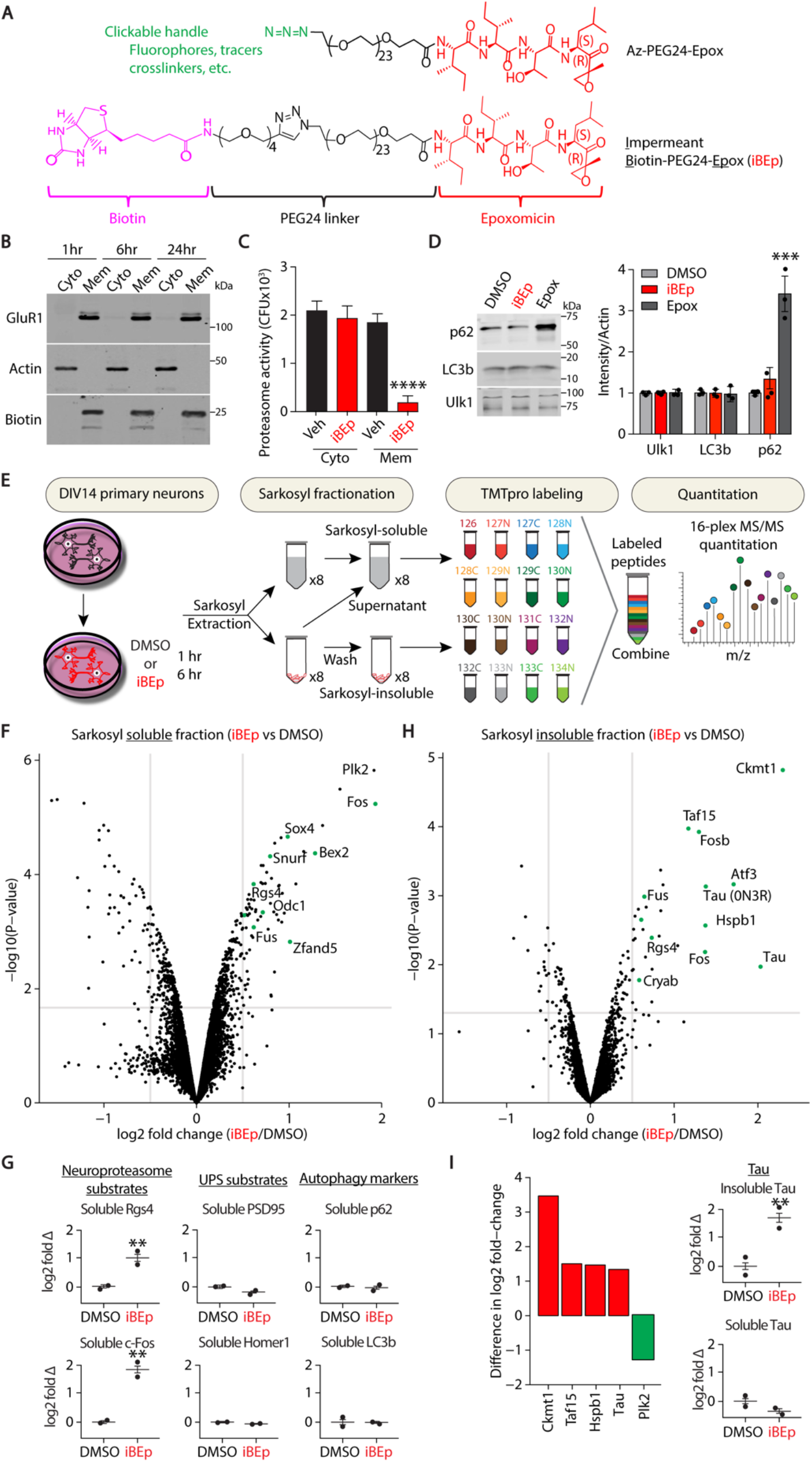
Quantitative proteomics reveals that selective inhibition of neuroproteasomes induces accumulation of sarkosyl-insoluble Tau. **(A)** (Top) Structure of azide-PEG24-epoxomicin handle to enable rapid conjugation of linkers to global proteasome inhibitor epoxomicin using Click chemistry. (Bottom) Structure of Neuroproteasome-specific inhibitor impermeant <u>B</u>iotin-PEG24-<u>Ep</u>oxomicin (iBEp). **(B)** Assessing permeability of iBEp in primary neurons. Primary cortical neurons obtained from WT mice were treated with 1 µM iBEp for 1, 6 and 24 hours and subjected to membrane fractionation. Cytosolic (Cyto) and membrane (Mem) fractions were immunoblotted using indicated antibodies. **(C)** Proteasome catalytic activity of membrane (Mem) and cytosolic (Cyto) fractions of neurons treated with iBEp. DIV14 primary cortical neurons obtained from WT mice were treated with DMSO (black) and iBEp (red) for 6 hours and subjected to membrane fractionation. Proteasome catalytic activity was assessed by monitoring degradation of Suc-LLVY-AMC; proteasome-dependent cleavage of the substrate increases AMC fluorescence. N=3 biological replicates. ****p<0.0001 by One-Way ANOVA. **(D)** Primary neurons treated with iBEp probed for autophagy markers. DIV14 primary cortical neurons obtained from WT mice were treated with DMSO (light gray), iBEp (red) or Epoxomicin (Epox, dark gray). Lysates were immunoblotted using indicated antibodies against common autophagy markers. Quantification of p62, LC3b, and Ulk1 intensities normalized to Actin. Data (right) are mean ± SEM normalized to respective DMSO controls. N=4 biological replicates, ***p<0.001 by One-Way ANOVA. **(E)** Schematic to analyze changes in sarkosyl-soluble and insoluble proteomes in response to neuroproteasome inhibition. DIV14 primary neurons obtained from WT mice were treated with DMSO or iBEp for 1 and 6 hours and subjected to sarkosyl fractionation. Sarkosyl-soluble and insoluble fractions were labeled with Tandem Mass Tags (TMTpro) for quantitative mass spectrometry. **(F)** Differential enrichment of proteins in the sarkosyl-soluble fraction of neurons treated with iBEp versus DMSO was analyzed and plotted as log2-fold change vs –log10(P-value). Green dots represent proteins enriched in the iBEp treatment compared to DMSO treatment. **(G)** Graphs plotting the log2-fold change of select Neuroproteasome substrates enriched in the sarkosyl-soluble fraction from MS analysis in (f) compared to UPS substrates and autophagy markers **p<0.01 (adjusted p-value after multiple corrections). **(H)** Differential enrichment of proteins in the sarkosyl-insoluble fraction of neurons treated with iBEp versus DMSO was analyzed and plotted as log2-fold change vs –log10(P-value). Green dots represent proteins enriched in the iBEp treatment compared to DMSO treatment. **(I)** (Left) Identification of proteins which are exclusively enriched in the insoluble fraction with no corresponding change in the soluble fraction (red). Plk2 represents the only protein which is enriched in the soluble fraction with no corresponding change in the insoluble fraction (green). Proteins plotted as differences in log2 fold changes. (Right) Graphs plotting the log2 fold changes of insoluble Tau (top) and soluble Tau (bottom) enriched in neurons treated with iBEp vs DMSO. **p<0.01 (adjusted p-value after multiple corrections).

We next tested the potency and selectivity of iBEp relative to Epoxomicin. Epoxomicin covalently and irreversibly binds the catalytic β subunits of the proteasome – therefore, the biotin moiety on iBEp serves as a tracer to detect whether proteasomes are inhibited by monitoring biotin signal on an SDS-PAGE gel(*83, 87-89*). We detect covalent biotinylation of only three proteins from iBEp-treated neurons, corresponding to the three catalytic subunits of the proteasome (**Fig S4F**). Therefore, we demonstrate that the iBEp linker does not modify the high specificity of iBEp compared to Epoxomicin alone. This is supported by previous data that modification of the non-reactive end of epoxomicin has been shown not to change the selectivity of Epoxomicin for the proteasome(*83, 87-89*). While iBEp is still highly active against the proteasome, we do find that the Biotin-PEG24 linker moderately reduces the efficacy of iBEp relative to Epoxomicin (**Fig S4G**). We therefore chose doses of inhibitor where the effective concentration of the inhibitor was similar to previous studies using Epoxomicin alone and well within the acceptable range of efficacy given how potent Epoxomicin is as a proteasome inhibitor.

To test the permeability of our compounds, we incubated primary cortical neurons with iBEp over a time course and prepared both cytosolic and plasma membrane-enriched fractions. Samples were immunoblotted and probed for cytosolic marker Actin and membrane protein GluR1 to validate our fractionation (**Fig 4B**). We found specific biotin signal corresponding to covalent inhibition of the proteasome by iBEp only in the membrane fraction, and not in the cytosol, over a 24 hour incubation in primary neurons (**Fig 4B**). As an orthogonal measure of cell penetrance, we tested whether the activity of cytosolic proteasomes was affected by iBEp treatment. We subjected cytosolic and membrane fractions from iBEp-treated neurons to LLVY-based proteasome activity assays. Cytosolic proteasome activity remains completely intact from iBEp-treated neurons compared to DMSO controls, but proteasome activity in the membrane fraction is almost entirely abrogated (**Fig 4C**). This suggests that neuroproteasomes are inhibited following iBEp incubation but cytosolic proteasomes remain unaffected.

To further validate our neuroproteasome-specific inhibitors, we sought a cellular functional readout of whether our cell-impermeant inhibitors affected cytosolic proteasomes. One of the most penetrant and robust consequences of total proteasome inhibition is the induction of autophagy as a cellular response mechanism to compensate for the lack of proteasomal degradation(*31-33, 90-92*). To test this, we assayed three autophagy markers, p62 levels, LC3bI/II ratios, and Ulk1 levels. Each are robust markers of different steps of autophagy. p62 has been demonstrated to be rapidly induced upon proteasome inhibition and LC3bI/II induction is thought to be slower, while Ulk1 is important for early phagosome formation(*93*). Cytosolic proteasome inhibition has been reproducibly demonstrated to induce a strong increase in p62 levels in neurons(*31, 32, 34, 93*). Consistent with these reported observations, we find a strong induction of p62 and little to no change in LC3bI/II following 12 hours of total proteasome inhibition with epoxomicin (**Fig 4D**). However, when we treat primary neurons with iBEp, we observe no change in either p62 or LC3bI/II levels (**Fig 4D**). These data are a functional demonstration that iBEp is cell-impermeable to neurons and signify that specifically inhibiting neuroproteasomes over all proteasomes does not induce compensatory autophagic degradation, and together, validate the use of iBEp as a neuroproteasome-specific inhibitor.

One important advantage of iBEp is the ability to use over extended timecourses, a property necessary to understand how neuroproteasomes shape the proteome, if at all. We hypothesized that prolonged neuroproteasome inhibition may reveal not only the turnover of candidate substrates, but also affect the solubility and aggregation of proteins. If true, this observation may provide some explanatory power for how neuroproteasome dysfunction (driven by ApoE4 or other mechanisms) could underlie deficits in proteostasis with potential relevance for neurodegenerative disorders. We incubated DIV12-14 primary cortical neurons with cell-impermeable proteasome inhibitors in biological duplicate for 1 and 6 hours, prepared sarkosyl-soluble and sarkosyl-insoluble samples, and then collected and processed samples for quantitative TMT-based proteomics (**Fig 4E**). We first analyzed proteins which increase over the time-course in the soluble fraction. We recapitulate our previously published data and find that neuroproteasome inhibition induces the accumulation of previously reported neuroproteasome substrates such as Rgs4, c-Fos, Sox4, Snurf, Arc, and Bex2 in the soluble fraction (**Fig 4F, G, Table S3**)(*4*). Importantly, we also do not see accumulation of classical ubiquitin-proteasome substrates such as PSD-95, Homer1, or GKAP(*94-96*) (**Fig 4G**). We also fail to observe an increase in p62, LC3b, GabarapL1, or any other marker of autophagy(*1, 97*) (**Fig 4G**). This provides orthogonal evidence that iBEp does not detectably penetrate cells and demonstrates the validity of this approach to determine the consequences of selective neuroproteasome inhibition. These validate our previous data identifying neuroproteasome substrates(*4*) and extend them by demonstrating neuroproteasome-dependent substrate turnover in a context without modulating neuronal activity and over extended periods of time.

Next, our quantitative proteomics analysis revealed a selective pattern of protein insolubility following neuroproteasome inhibition. We find that only 52 proteins become sarkosyl-insoluble in response to neuroproteasome inhibition (**Fig 4H, Table S3**), in contrast with over 250 proteins increasing in the soluble fraction (**Fig 4F**). To narrow in on this list, we quantitated which proteins show a greater than log-fold change in the insoluble fraction, but showed no change in the soluble fraction (**Fig 4I**). This analysis identified only four proteins which show significant increases in the insoluble fraction relative to the soluble fraction (Ckmt1, Tau, Taf15, and Hspb1) and one protein which significantly increases in the soluble fraction and showed no change in the insoluble fraction (Plk2) (**Fig 4I**). Of note, Taf15 has previously been observed in detergent-insoluble fractions of brains from Alzheimer’s patients, correlated with both Amyloid β and Tau insolubility(*98-101*). Ckmt1 has also been found in Neurofibrillary tangles and interacting with phosphorylated Tau inclusions(*102-104*).

### Neuroproteasomes are differentially regulated by ApoE isoforms to determine formation of endogenous sarkosyl-insoluble Tau inclusions

Our findings raise the intriguing possibility that a fraction of endogenous mouse Tau becomes sarkosyl-insoluble following neuroproteasome inhibition. We sought to validate these findings by immunoblot and extend them to determine if the same principle would hold for human Tau isoforms, without pathogenic and aggregation-driving mutations. We therefore cultured primary mouse cortical neurons from either WT or hTau-KI mice (*105*). The entire mouse *Mapt* sequence is replaced with the human *Mapt* sequence (including upstream gene regulatory elements) at the endogenous locus in hTau-KI mice. hTau-KI mice express all six human isoforms of Tau protein using the endogenous *Mapt* regulatory elements and therefore, hTau is expressed at normal physiological levels(*105*). DIV12-14 neurons were treated with either the neuroproteasome inhibitor iBEp or the cell-permeable proteasome inhibitor Epoxomicin. We then performed sarkosyl fractionations to separate treated neurons into soluble and insoluble fractions and immunoblotted these fractions for total Tau and GAPDH. The R2295 antibody reacts against both total human and total mouse Tau and is highly specific, making it the ideal tool to monitor endogenous Tau in WT and hTau-KI samples(*106*). We observed no Tau signal in primary neurons cultured from Tau knockout (Tau-KO) mice, demonstrating the specificity of our antibodies and approach (**Fig S5A**)(*106*). We found a significant increase in sarkosyl-insoluble Tau from both the WT (**Fig 5A**) and hTau-KI neurons treated with iBEp (**Fig 5B, S5B**). Intriguingly, we find no change in sarkosyl-insoluble Tau in neurons treated with the cell-permeable Epoxomicin compared to DMSO controls, despite the fact that Epoxomicin inhibits cytosolic proteasomes and neuroproteasomes (**Fig 5A,B**). We reason that Epoxomicin inhibits neuroproteasomes and likely induces the formation of Tau inclusions (similar to iBEp), but these Tau inclusions cannot be detected because simultaneous inhibition of cytosolic proteasomes induces autophagy and other compensatory mechanisms which can clear these inclusions. This claim is supported by an extensive body of work on Tau clearance mechanisms(*31-35, 93*). We posit that such compensatory mechanisms are not triggered when neuroproteasomes are specifically inhibited, which is consistent with our observations that Epoxomicin induces p62 expression whereas iBEp does not (**Fig 4D, G**).

**Figure 5:**
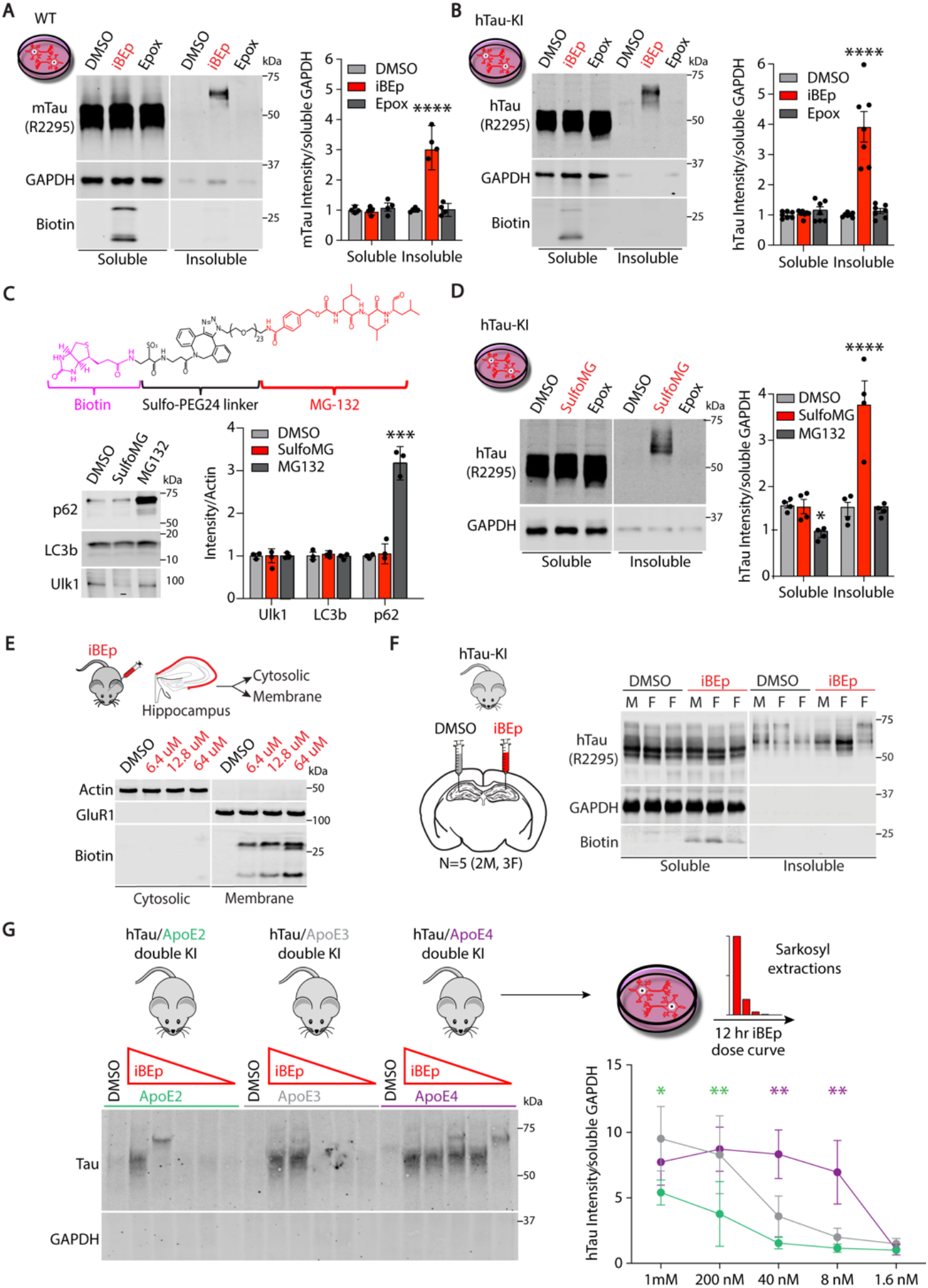
Neuroproteasomes are differentially regulated by ApoE isoforms to determine formation of endogenous sarkosyl-insoluble Tau inclusions. **(A)** Effect of neuroproteasome-specific inhibitor iBEp on Tau aggregation in primary neurons. DIV14 primary cortical neurons obtained from WT mice were treated with DMSO (light gray), 1µM iBEp (red), or 1 µM Epoxomicin (Epox, dark gray) for 12 hours and were subjected to sarkosyl fractionation. Sarkosyl-soluble (Soluble) and sarkosyl-insoluble (Insoluble) fractions were immunoblotted using following antibodies: R2295 for total Tau, GAPDH, and fluorescent Streptavidin (Biotin). Quantification of soluble and insoluble murine Tau (mTau) intensities normalized to soluble GAPDH. Data (right) are mean ± SEM normalized to respective DMSO controls. N=4 biological replicates, ****p<0.0001 by One-Way ANOVA Tukey’s Multiple Comparison Test, sarkosyl-soluble and insoluble analyzed separately. **(B)** Effect of neuroproteasome inhibition on aggregation of human Tau (hTau). Experiment done identically as in (a) but using DIV14 primary cortical neurons obtained from hTau-knock-in (hTau-KI) mice instead of WT mice. Data (right) are mean ± SEM normalized to respective DMSO controls, N=7 biological replicates, ****p<0.0001 by One-Way ANOVA Tukey’s Multiple Comparison Test. **(C)** Generation of reversible neuroproteasome inhibitors to test effects of neuroproteasome inhibition. (Top) Structure of Sulfo-MG132 (SulfoMG). (Bottom) DIV14 hTau-KI primary cortical neurons were treated with DMSO (light gray), 1µM SulfoMG (red), or 1µM MG132 (dark gray), with or without 10µg/mL cycloheximide (CHX), and immunoblotted using indicated antibodies. Quantification of p62, LC3b, Ulk1 intensities normalized to Actin. -CHX data (right) are mean ± SEM normalized to DMSO. N=3 biological replicates, ***p<0.001 by One-Way ANOVA Tukey’s Multiple Comparison Test. **(D)** Effect of reversible neuroproteasome inhibition using SulfoMG on Tau aggregation. DIV14 primary cortical neurons obtained from hTau-KI mice were treated as described in (e) and processd as described in (b). Sarkosyl-soluble (Soluble) and sarkosyl-insoluble (Insoluble) fractions were immunoblotted using indicated antibodies and fluorescent Streptavidin (Biotin). Quantification of soluble and insoluble Tau intensities were normalized to soluble GAPDH. Data (right) are mean ± SEM normalized respective to DMSO controls. N=4 biological replicates, *p<0.05, ****p<0.0001 by One-Way ANOVA Tukey’s Multiple Comparison Test. **(E)** Testing iBEp permeability *in vivo*. (Top) iBEp or 0.7% DMSO was stereotactically injected into the CA1 region of the hippocampus in the left hemisphere of WT mice. 72 hours post injection, hippocampi were isolated and subjected to membrane fractionation without detergent. (Bottom) Cytosolic and membrane fractions were immunoblotted using indicated antibodies and fluorescent Streptavidin (Biotin). **(F)** Effect of neuroproteasome inhibition on Tau aggregation *in vivo*. Four-to-five month old hTau-KI mice were stereotactically injected with iBEp (red) in the CA1 region of the left hippocampus and DMSO (gray) was injected contralaterally. The hippocampi were subjected to sarkosyl fractionation 72 hours post injection. The sarkosyl-soluble (Soluble) and sarkosyl-insoluble (Insoluble) fractions were immunoblotted using indicated antibodies and fluorescent streptavidin (Biotin). Quantification of soluble and insoluble Tau intensities normalized to soluble GAPDH. Data shown (right) are mean ± SEM normalized to DMSO. N= 5 biological replicates; (2M, 3F), *p<0.05, ***p<0.001 by One-Way ANOVA Tukey’s Multiple Comparison Test. **(G)** Effect of ApoE isoforms on neuroproteasome-dependent Tau aggregation. DIV14 primary neurons from hTau/ApoE2 double KI (green), hTau/ApoE3 double KI (gray), and hTau/ApoE4 double KI (purple) mice were incubated with the following iBEp (red) doses: 1000, 200, 40, 8 and 1.6nM. Treated neurons were subjected to sarkosyl extraction and sarkosyl-insoluble fractions were immunoblotted using indicated antibodies. Quantification of sarkosyl-insoluble Tau intensity was normalized to soluble GAPDH. Data are mean ± SEM normalized to respective DMSO controls. N=3 biological replicates, **p<0.01, *p<0.05 by Two-Way ANOVA Tukey’s Multiple Comparisons Test.

Many of our experiments rely on the use of iBEp because the biotin tracer on an irreversible and specific inhibitor allows us to track which proteasomes are inhibited. However, we wanted to validate our findings using a proteasome inhibitor with a distinct mechanism of action from Epoxomicin. We therefore modified MG132, a reversible peptide aldehyde proteasome inhibitor, with a sulfonated biotin linker on the nonreactive N-terminus(*84*). This negatively charged water-soluble linker renders Sulfo-MG132 (SulfoMG) cell-impermeable, much like the historical use of sulfonated linkers to perform surface biotinylation (**Fig 5C).** We observe efficient conjugation by HPLC analysis (**Fig S5C, D**). All molecules were HPLC-purified as single peaks and confirmed by LC/MS analysis (**Fig S5E, F**). We find significant inhibition of proteasomes using SulfoMG despite the moderate loss of efficacy compared to MG132 as a consequence of the linker (**Fig S5G**). MG132 is a reversible inhibitor and therefore, we cannot detect the covalent modification of the proteasome subunits. Instead, we turned to cellular and functional readouts of permeability. We found a similar induction of autophagy markers in neurons treated with MG132 as with epoxomicin and a similar lack of induction with SulfoMG as with iBEp (**Fig 5C)**. Confident in its use as a chemically orthogonal cell-impermeable proteasome inhibitor, we next incubated DIV13 primary hTau-KI neurons with SulfoMG. We find that SulfoMG induces similar partitioning of hTau into the sarkosyl-insoluble fraction as iBEp (**Fig 5D**). Moreover, we find that MG132, like Epoxomicin, does not affect the sarkosyl-insoluble accumulation of Tau (**Fig 5D**).

Next, we examined whether neuroproteasome inhibition would induce sarkosyl-insoluble endogenous Tau inclusions *in vivo*. Here, we only used iBEp and not SulfoMG for three reasons: first, because SulfoMG is reversible and is therefore subject to being washed out by CSF flow *in vivo*; second, because we could biochemically monitor iBEp permeability using the presence of the biotin signal; and third, because Epoxomicin is a more specific proteasome inhibitor than MG132. We tested iBEp permeability *in vivo* by stereotactically injecting iBEp into the CA1 region of the mouse hippocampus for 72 hours. Hippocampi were microdissected and separated into cytosolic and membrane-enriched fractions and samples were immunoblotted (**Fig 5E**). We fail to observe penetration into the cytosol until we inject 64μM, so we chose to inject at 10-fold under this dose as our maximal concentration (**Fig 5E**). This *in vivo* dose is consistent with and far lower than previous studies using Epoxomicin alone (9.0 and 13.5mM (*107, 108*)). We performed stereotactic injections of 6.4 uM iBEp and contralateral injections of the vehicle (0.07% DMSO) into the hippocampus of hTau-KI and Tau-KO mice. After 72 hours, we collected the hippocampi and performed sarkosyl extractions into soluble and insoluble fractions. We fail to observe any detectable Tau in the Tau-KO mice in DMSO- or iBEp-injected hippocampi (**Fig S5H**). Mirroring our data in primary neurons, we find a strong increase of sarkosyl-insoluble Tau in hippocampi from hTau-KI mice injected with iBEp (**Fig 5F**).

In our experiments, we find that inhibiting neuroproteasomes to saturation induces the accumulation of sarkosyl-insoluble Tau. Taken together with our data demonstrating that ApoE isoforms regulate neuroproteasome localization (ApoE4<E3<E2), we predicted that neurons from ApoE4-KI mice would be more susceptible accumulating sarkosyl-insoluble Tau relative to neurons from ApoE3-KI or ApoE2-KI mice. To test this prediction, we crossed each of the ApoE-KI lines to the hTau-KI lines, generating mice which endogenously express ApoE2, E3, or E4 as well as hTau. DIV14 primary neurons from each line (hTau/ApoE2 double KI, hTau/ApoE3 double KI, and hTau/ApoE4 double KI) were then treated with a range of iBEp doses spanning four log units. Neurons were then separated into sarkosyl-soluble and insoluble fractions and immunoblotted. We found no sarkosyl-insoluble Tau aggregates at baseline in any line, which suggests that the ApoE4-induced reduction of neuroproteasomes levels alone is not sufficient to drive Tau aggregation (**Fig 5G, S5I**). However, after treatment with even the lowest dose of iBEp, we found accumulation of sarkosyl-insoluble Tau in hTau/E4-dKI neurons, rendering E4 neurons over 25-fold more susceptible to Tau aggregation than E3 neurons (**Fig 5G**). Conversely, neurons from hTau/E2-dKI mice were over 10-fold less susceptible to neuroproteasome-inhibition induced Tau aggregation than neurons from hTau/E3-dKI mice (**Fig 5G**). Overall, these data demonstrate that ApoE isoforms differentially shift the threshold for the neuroproteasome-dependent formation of endogenous sarkosyl-insoluble Tau inclusions.

### Neuroproteasome inhibition induces endogenous phosphorylated, sarkosyl-insoluble, Thioflavin S-positive Tau aggregates

We noticed that neuroproteasome inhibition induces the shift of sarkosyl-insoluble Tau into a species which migrates at a higher molecular weight on an SDS-PAGE gel, at approximately 64kDa (**Fig 5A, B, D, F, G, S5B, S6A)**. Similar shifts in molecular weight of Tau are observed in the AD brain and is a hallmark phenotype of multiple neurodegenerative disorders (*109, 110*). This molecular weight shift can in part be due to Tau phosphorylation at many residues. Therefore, we wanted to address whether iBEp-induced Tau aggregates are phosphorylated. Rather than profiling by a panel of phospho-specific antibodies against Tau, we took a more unbiased approach and measured Tau phosphorylation by quantitative phosphoproteomics. We treated primary WT neurons with iBEp for either 1, 6, or 24 hours and then performed sarkosyl fractionations. Soluble and insoluble fractions were processed for TMT labelling in preparation for quantitative mass spectrometry and then peptides were run over an Fe-NTA column to enrich for phospho-peptides (**Fig 6A**). We then used this enriched population for quantitative phospho-proteomics analysis at the single peptide level (**Table S4**). Here, we find a time-dependent increase in Tau phosphorylation at only 4 of 26 identified sites which correspond to human S202, T205, T217, and S404 (**Fig 6A**). This is a specific signature of phosphorylation which corresponds to sites at which Tau is phosphorylated in the AD brain and detected by established antibodies (S202 – CP13(*111*), T205 – AT8(*112*), and S404 – PHF1(*113*)) as well as CSF biomarkers for AD progression (T217)(*114*).

**Figure 6:**
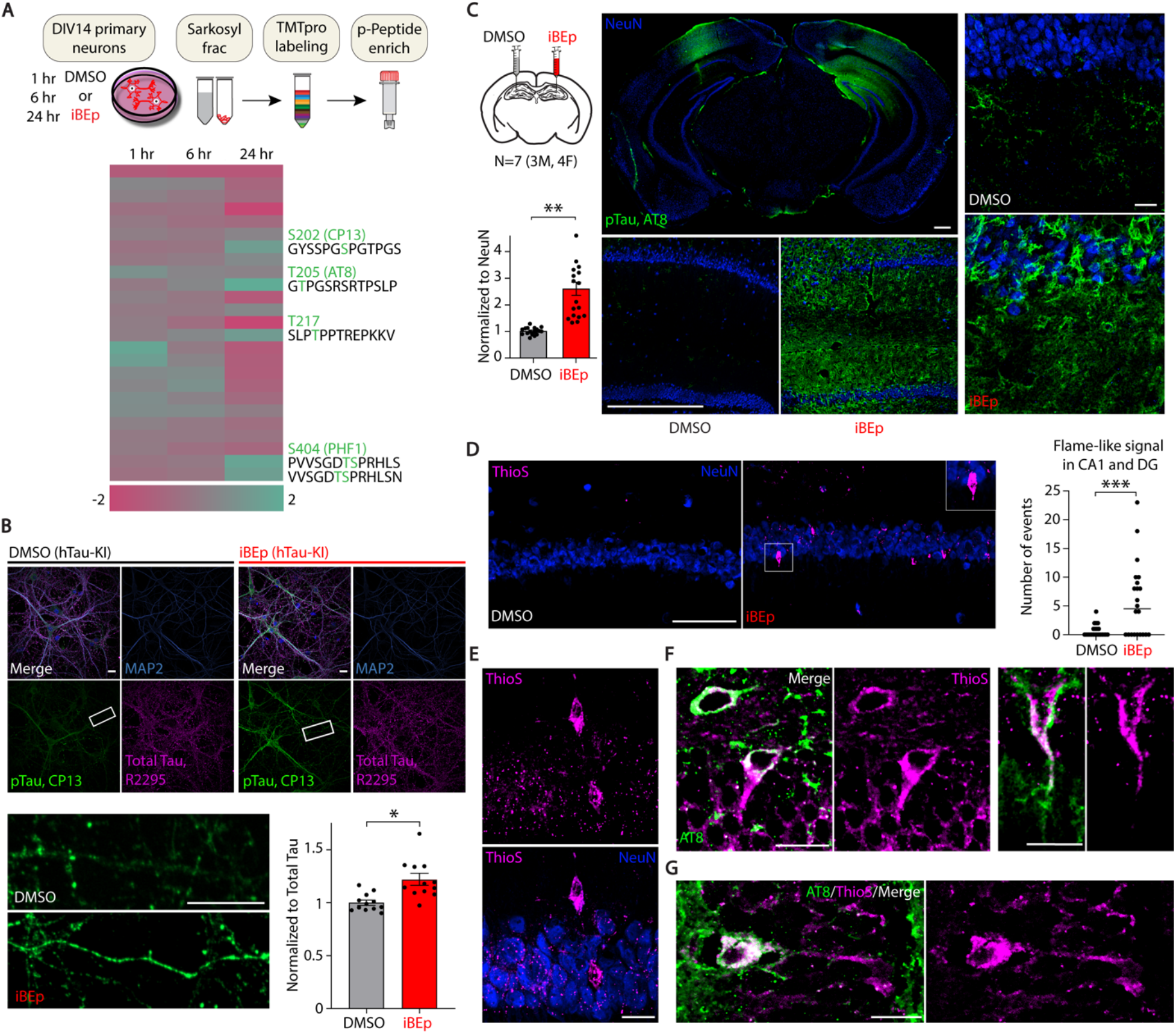
Neuroproteasome inhibition induces endogenous phosphorylated, sarkosyl-insoluble, Thioflavin S-positive Tau aggregates. **(A)** (Top) Schematic of quantitative phosphoproteomics experiments from primary neurons treated with neuroproteasome inhibitor iBEp. DIV14 primary neurons obtained from WT mice were treated with 1µM iBEp for 1, 6, and 24 hours or DMSO and separated into sarkosyl soluble or insoluble (red) fractions. Samples were processed for quantitative TMTpro-based phosphoproteomic analysis. (Bottom) Heatmap displays all identified Tau phosphopeptides and their relative depletion (pink) or enrichment (green) compared to DMSO controls at indicated timepoints. Select phosphopeptides enriched in iBEp treated neurons over the timecourse are labeled (green) and antibodies to detect these phosphoepitopes indicated in parenthesis. **(B)** Immunocytochemical analysis of neurons treated with iBEp to measure Tau phosphorylation. DIV14 primary hippocampal neurons obtained from hTau-knock-in (hTau-KI) mice treated with DMSO or iBEp (red) for 12 hours were fixed and stained using indicated antibodies: MAP2 (blue) to label dendrites, phosphorylated Tau (pTau, CP13, green) or total Tau (R2295, purple). Quantification of pTau signal intensity was normalized to total Tau. Data are mean ± SEM normalized to DMSO. N=12 (3 biological replicates, 4 images/replicate), *p < 0.05 by Paired T-test. Scale bars=15µm **(C)** Immunohistochemical analysis of mice stereotactically injected with iBEp to measure Tau phosphorylation *in vivo*. Four-to-five month old hTau-KI mice were stereotactically injected (schematic) with iBEp (red) into the CA1 region of the hippocampus in the left hemisphere and DMSO (gray) was injected contralaterally (N=7 (3M, 4F)). Mice were collected 72 hours post injection and sections were stained using indicated antibodies: NeuN (blue) and AT8 (pTau, green). 4X, 20X and 60X representative images are shown. Quantification of pTau signal intensity was normalized to NeuN intensity. Analysis was done blinded to experimental condition. Data are mean ± SEM normalized either to DMSO or iBEP+CHX. n=2 sections/animal, **p<0.01 by Paired T-test/experiment. Scale bars=500µm (left), scale bar=25µm (right). **(D)** Immunohistochemical analysis of mice stereotactically injected with iBEp to measure Thioflavin-S positive Tau aggregates *in vivo*. Four-to-five month old hTau-KI mice were treated identically to (c), but stained for Thioflavin S (magenta) to visualize β-sheet containing aggregates and immunostained with NeuN (blue). N=7 (3M, 4F). Left, Vehicle-treated hippocampus, Right, iBEp-treated hippocampus (red). Inset in iBEp-treated hippocampus indicates representative example of flame-like Thioflavin-S positive inclusion. Data are quantification of number of flame-like Thioflavin-S positive inclusions. Counting and analysis was blinded to experimental condition. ***p<0.001 by one-way ANOVA. Scale bars=100µm. **(E)** Higher magnification image of flame-like Thioflavin S-positive inclusions (magenta), NeuN (blue). Top, ThioS alone, Bottom, merge with NeuN. Scale bar=20µm. **(F, G)** Hippocampal sections from iBEp-injected mice co-stained with Thioflavin S (magenta) and AT8 (Green), overlap appears white. Images are single Z-plane images to accurately demonstrate co-localization of ThioS and AT8 staining, demonstrating that some ThioS+ inclusions co-localize with AT8. (F) Contains two representative examples of flame-like ThioS+ inclusions while (G) displays representative example of thread-like inclusions. Scale bar=20µm.

To validate these data, we treated hTau-KI neurons with iBEp and then fixed and stained these neurons. We observed no staining in neurons cultured from Tau-KO mice using the R2295 antibody to detect total Tau or the CP13 antibody to detect phosphorylated S202 (**Fig S6B**). Incubating neurons with secondary antibodies alone showed no signal (**Fig S6C**). We therefore exclusively used CP13 for our ICC experiments to detect phosphorylated Tau. We find a significant elevation of CP13 signal in iBEp-treated primary hippocampal hTau-KI neurons compared to controls (**Fig 6B**). This iBEP-induced signal appears punctate and distributed in both axons and MAP2+ dendrites (**Fig 6B**). To validate these data, we conducted immunohistochemical (IHC) analysis of sections from hTau-KI mice stereotactically injected with iBEp and contralaterally injected with vehicle control (0.07% DMSO). After stereotactic injections, animals were allowed to recover for 72 hours and then were perfused and sectioned for IHC. We observed no staining in the Tau-KO mice using the AT8 antibody or with secondary antibodies alone (**Fig S6D**, **S6E**). We observe a threefold induction of phosphorylated Tau signal in hippocampi injected with iBEp compared to the contralateral PBS controls. We see a strong increase in AT8+ inclusions mislocalized to somatodendritic compartments in iBEp-injected hippocampi, which extend into the stratum radiatum and stratum moleculare (**Fig 6C**).

While sarkosyl-insoluble Tau reflects aggregated Tau species(*99, 106, 115-117*), there are varying definitions of a protein aggregate: misfolded proteins which cluster together, proteins which form stable inclusions, inclusions which are resistant to varying detergents, and then in some cases, proteins which form very stable aggregates and even amyloids(*118, 119*). Thioflavin S (ThioS) is a dye that stains the β-sheet structures of amyloid aggregates, including both aggregated Tau tangles and amyloid plaques. Therefore, ThioS is used for the neuropathological diagnosis of a wide variety of neurodegenerative disorders(*8, 54, 120*). We conducted ThioS staining of sections from hTau-KI mice stereotactically injected with iBEp and contralaterally injected with vehicle control. After stereotactic injections, animals were allowed to recover for 72 hours and then brains were perfused, sectioned, and stained for ThioS. We find a strong increase in ThioS staining from the CA1 region of hippocampi exposed to iBEp relative to vehicle controls (**Fig 6D**). ThioS+ inclusions appear as flame-like inclusions in both the CA1 pyramidal layer and the granule cell layer of the Dentate Gyrus (**Fig 6D, E, S6F-J**). Neuroproteasome inhibition-induced flame-like ThioS+ aggregates co-localize with both phosphorylated Tau (AT8) and total Tau (DA9) (**Fig 6F, G, S6K**) in single Z-plane images from neurons in the CA1 and DG of hippocampus. We conclude that some neuroproteasome-induced aggregates we observe are indeed composed of Tau. In addition to these large ThioS+ inclusions, we also observe some thread-like ThioS+ inclusions in the hippocampal neuropil as well as widely distributed thin ThioS+ inclusions that appear dispersely throughout the iBEp-injected hippocampus (**Fig 6G, S6L, M**). The ThioS+ inclusions formed following neuroproteasome inhibition appear to resemble neurofibrillary tangle (NFT)-like and neuropil thread-like inclusions observed in various Tauopathies including AD (*8, 17, 54, 120*). While we find it highly surprising that ThioS+ Tau inclusions appear just 72 hours after neuroproteasome inhibition, these data support our overall conclusion that neuroproteasome inhibition induces the aggregation of endogenous Tau.

### *De novo* protein synthesis is required for neuroproteasome-dependent induction of endogenous Tau aggregates

Neuroproteasomes are 20S complexes and do not have the machinery to recognize ubiquitin or unfold substrates(*3, 4*). Instead, neuroproteasomes co- or peri-translationally degrade newly synthesized and unfolded substrates rather than substrates which are fully folded(*4*). We therefore tested whether new protein synthesis was required for the neuroproteasome-dependent formation of endogenous Tau aggregates. We incubated primary neuronal cultures with either iBEp or Epoxomicin and then co-incubated these cultures with cycloheximide, which blocks translation elongation. Here, we reproduce iBEp-induced Tau aggregation and find that cycloheximide blocks this iBEp-mediated increase in both WT primary neurons (**Fig 7A**) and in hTau-KI primary neurons (**Fig 7B**). Cycloheximide also blocks the effect of SulfoMG on the formation of endogenous insoluble Tau inclusions, consistent with our observations using iBEp (**Fig 7C**). Next, we tested whether inhibition of protein synthesis blocks the accumulation of sarkosyl-insoluble Tau *in vivo* in hTau-KI mice. We stereotactically co-injected cycloheximide and iBEp together contralateral to iBEp alone into the hippocampus. We find a dramatic decrease in sarkosyl-insoluble Tau in the hippocampus from mice co-injected with iBEp and cycloheximide compared to those injected with iBEp alone (**Fig 7D, S7A**). Next, co-injection of cycloheximide with iBEp completely eliminated AT8 staining compared to contralateral injection of iBEp alone (**Fig 7E**). Finally, similar to what we observed with sarkosyl-insoluble Tau and AT8 staining, we also observe a complete return to baseline of ThioS+ staining in hippocampi co-injected with iBEp and cycloheximide relative to those injected with iBEp alone (**Fig 7F**).

**Figure 7.**
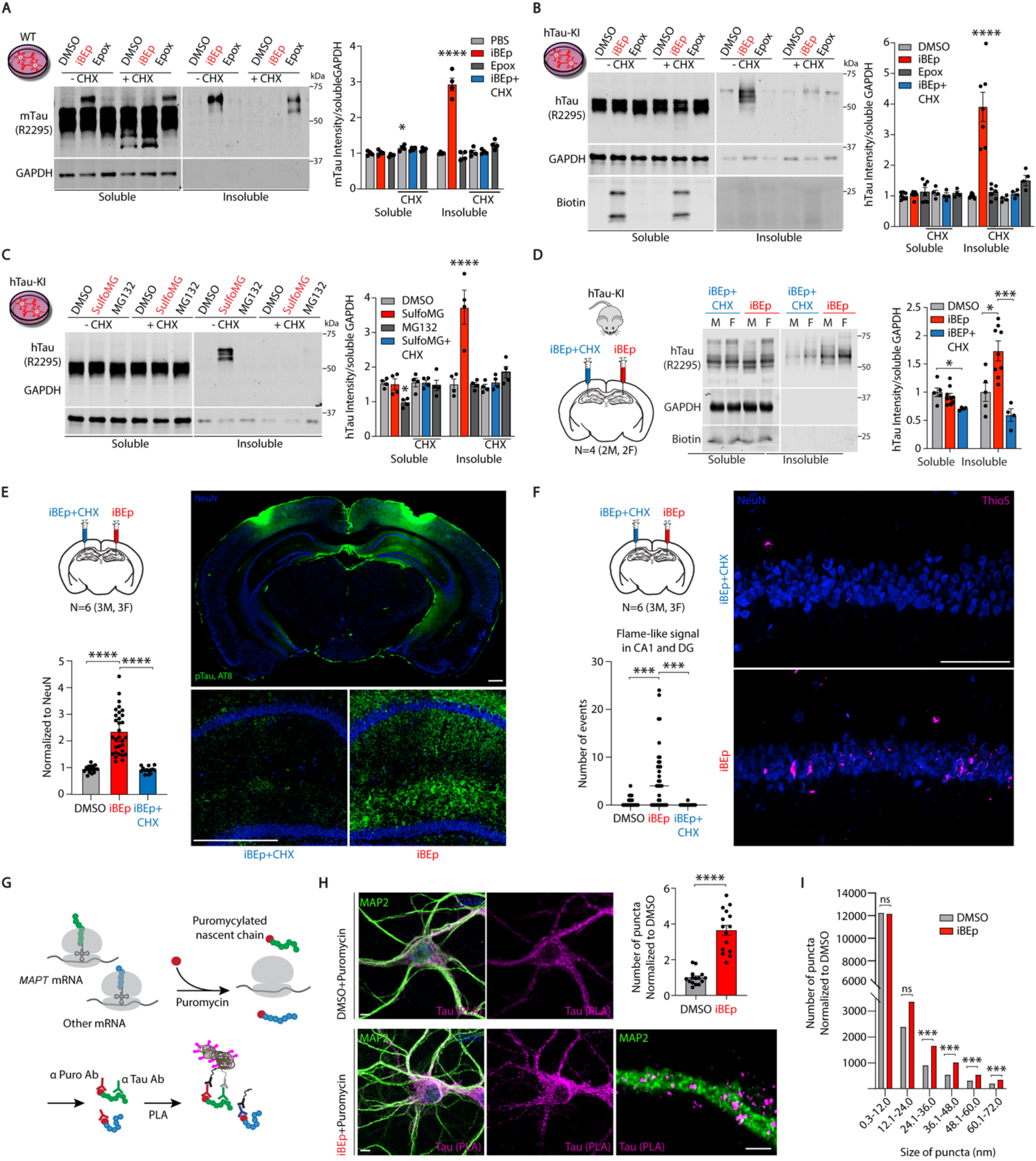
*De novo* protein synthesis is required for neuroproteasome-dependent induction of endogenous Tau aggregates. **(A)** Effect of protein synthesis inhibition on neuroproteasome-dependent Tau aggregation. DIV14 primary cortical neurons obtained from WT mice were treated with DMSO, 1µM iBEp, or 1 µM Epoxomicin (Epox) for 12 hours, with and without cycloheximide (CHX). Sarkosyl-soluble (Soluble) and sarkosyl-insoluble (Insoluble) fractions were immunoblotted using following antibodies: R2295 for total Tau, GAPDH, and fluorescent Streptavidin (Biotin). iBEp + CHX indicated in blue for emphasis relative to iBEp alone (red). Data (right) are mean ± SEM normalized to respective DMSO controls. N=4 biological replicates, ****p<0.001 by One-Way ANOVA Tukey’s Multiple Comparison Test. **(B)** Effect of protein synthesis inhibition on neuroproteasome-dependent Tau aggregation. DIV14 primary cortical neurons obtained from hTau-KI mice were treated and processed identically to (a). iBEp + CHX indicated in blue for emphasis relative to iBEp alone (red). Data (right) are mean ± SEM normalized to respective DMSO controls. N=7 biological replicates without CHX, N=4 with CHX, ****p<0.001 by One-Way ANOVA Tukey’s Multiple Comparison Test. **(C)** Effect of protein synthesis inhibition on neuroproteasome-dependent Tau aggregation. DIV14 primary cortical neurons obtained from hTau-KI mice were treated with DMSO, 1µM SulfoMG, or 1 µM MG132 for 12 hours, with and without cycloheximide (CHX) and processed identically to (a). SulfoMG + CHX indicated in blue for emphasis relative to SulfoMG alone (red). Data (right) are mean ± SEM normalized to respective DMSO controls. N=4 biological replicates, ****p<0.001 by One-Way ANOVA Tukey’s Multiple Comparison Test. **(D)** Effect of protein synthesis inhibition on neuroproteasome-dependent Tau aggregation *in vivo*. Four-to-five month old hTau-KI mice were stereotactically injected with iBEp in the left hippocampus and iBEp + CHX were co-injected (blue) contralaterally. Hippocampi were subjected to sarkosyl fractionation 72 hours post injection. The sarkosyl-soluble (Soluble) and sarkosyl-insoluble (Insoluble) fractions were immunoblotted using indicated antibodies and fluorescent streptavidin (Biotin). Quantification of soluble and insoluble Tau intensities normalized to soluble GAPDH. Data shown (right) are mean ± SEM normalized to DMSO. N=4 biological replicates (2M, 2F), *p<0.05, ***p<0.001 by One-Way ANOVA Tukey’s Multiple Comparison Test. **(E)** Four-to-five month old hTau-KI mice were stereotactically injected (schematic) with iBEp (red) and iBEp + CHX (blue) contralaterally (N=6, (3M, 3F)). Mice were collected 72 hours post injection and sections were stained using indicated antibodies: NeuN (blue) and AT8 (pTau, green). 4X, 20X and 60X representative images are shown. Quantification of pTau signal intensity was normalized to NeuN intensity. Analysis was done blinded to experimental condition. Data are mean ± SEM normalized to DMSO. n=3 sections/animal, ****p<0.0001 by one-way ANOVA. Scale bar=500µm **(F)** Immunohistochemical analysis of hTau-KI mice stereotactically injected (schematic) with iBEp (red) and iBEp + CHX (blue) contralaterally (N=6, (3M, 3F)). Mice were collected 72 hours post injection and sections and stained with Thioflavin-S and antibodies against NeuN. Thioflavin S–labeled β-sheet protein structure is stained in magenta, NeuN is stained in blue. Data are quantification of number of flame-like Thioflavin-S positive inclusions. Counting and analysis was blinded to experimental condition. ***p<0.001 by one-way ANOVA. Scale bar=100µm (right). **(G)** Schematic of puro-PLA-Tau experiment. Newly synthesized Tau (green) and other nascent proteins (blue) labeled with puromycin (red). Coincidence detection between anti-puromycin and anti-Tau antibodies result in an *in situ* ligation and amplification reaction which can be detected by fluorescent oligos (pink). **(H)** Neuroproteasome inhibition induces accumulation of newly synthesized Tau. Primary DIV14 hippocampal neurons treated with DMSO or iBEp (red) were puromycylated for 10 minutes. Tau-PLA-Puro labeling (pink) denotes newly synthesized Tau puncta, total Map2 positive dendrites (green), DAPI (blue). Higher magnification super-resolution image to bottom right. Quantification of Tau-PLA-Puro signal intensity was normalized to DMSO. ****p<0.0001 by Paired T-test. Scale bar= 5µm. **(I)** Size distribution of Tau-PLA-Puro puncta. Puncta were binned into varying sizes and then number of puncta were normalized to the number of puncta seen in each bin with DMSO treatment. This normalizes for the increase in puncta observed with iBEp treatment, allowing for a fair comparison in size. ***p<0.001 by Paired T-test/size bin.

There are many mechanisms which could explain our observation that new protein synthesis is required for the formation of neuroproteasome inhibition-induced endogenous Tau aggregates. We interpret our data and argue that neuroproteasome inhibition induces the accumulation of newly synthesized protein(s) important for Tau accumulation. The most straightforward candidate protein would be Tau itself. To determine specifically if we observed more newly synthesized Tau protein following neuroproteasome inhibition, we took advantage of proximity ligation assays (PLA) coupled with puromycylation to visualize newly synthesized Tau protein(*121, 122*). Briefly, we used specific monoclonal N-terminal antibodies against Tau and specific antibodies against puromycin. Each antibody is recognized by secondary antibodies tethered to oligonucleotides that can ligate and amplify *in situ*. When antibodies against Tau and Puromycin are proximal to each other, this permits the oligonucleotide ligation which can be detected by a fluorescent signal under hybridizing conditions (**Fig 7G**). A positive PLA signal therefore indicates the presence of newly synthesized Tau. We incubated Puromycin in primary neuronal cultures for only 10 minutes, which allows us to visualize a defined short snapshot of newly synthesized Tau. We observe no signal in neurons either not treated with puromycin or in neurons where we block protein synthesis using cycloheximide or anisomycin during puromycylation, validating our approach (**Fig S7B, C**). We observe a striking ∼3.5-fold increase in Puro-PLA-Tau signal in iBEp-treated neurons relative to DMSO controls (**Fig 7H**). We then analyzed the size distribution of diameters of the newly synthesized Tau puncta (**Fig 7I**). After accounting for and normalizing against the increase in puncta number seen in iBEp-treated neurons, we observe an increase in the size distribution of PLA puncta from iBEp treated neurons. We find that iBEp treated neurons contain an over twofold increase in newly synthesized Tau puncta greater than 24nm in diameter (**Fig 7I**). We interpret this to mean that there are larger inclusions of newly synthesized Tau which form following neuroproteasome inhibition, consistent with our observations that neuroproteasome inhibition induces Tau aggregation.

## Discussion

We conclude that neuroproteasome localization is differentially regulated by ApoE isoforms, which is critically important because reduced neuroproteasome function induces the aggregation of endogenous Tau. The relevance for neuroproteasomes in many aspects of the cell biology of neurons, as well as pathophysiology of neurodegeneration, will be critical to explore in the future. The cell-impermeable neuroproteasome-specific inhibitors we have developed and the tools we have built to visualize and purify neuroproteasomes will enable the broad study of neuroproteasomes in various aspects of neuronal physiology and pathophysiology.

The mechanisms by which ApoE interacts with and influences neuroproteasome localization remain unexplored. ApoE is found as a lipoprotein and the classic role for ApoE is a cholesterol carrier(*123*). While we demonstrate only lipidated ApoE can influence neuroproteasome localization, we do not understand whether the lipids transported by ApoE are involved in regulating neuroproteasomes or if the lipoproteins are more efficiently able to interact with the neuroproteasome complex. Neuroproteasome levels change in ApoE3/3 AD carriers relative to controls, which likely suggests that other mechanisms besides ApoE are at play in controlling neuroproteasome localization. It is therefore an open and important task to define the milieu of molecular and cell biological factors that can influence neuroproteasome localization. Detailed characterization of such mechanisms may help understand whether and how neuroproteasomes contribute to the large diversity of phenotypes in Tauopathies and neurodegenerative disease. Along these lines, ApoE has large effects on both amyloid deposition and Tau aggregation based on neuropathological data (*124*). Future work will determine whether and how neuroproteasomes can impinge on the amyloid pathway, if at all.

In the broadest sense, we find that neuroproteasome localization and function is a determinant for the proteostatic capacity of neurons to defend against protein aggregation. A major conclusion of our work is that neuroproteasome-specific inhibition induces the formation of endogenous Tau inclusions, suggesting the neuroproteasome is a pivotal mechanism in determining Tau aggregate formation. This function is notably distinct from the canonical role of proteasomes in protein clearance. Sarkosyl-insoluble Tau aggregates formed by neuroproteasome inhibition in neurons are endogenous, without a need for pathogenic mutations. We suggest that this may be important for Tau aggregation observed in a high percentage of sporadic AD patients who do not have genetic lesions in the *MAPT* locus. It will likely be important to determine how neuroproteasome inhibition-induced endogenous Tau aggregates compare to other types of misfolded Tau.

One of the key observations we make is that new protein synthesis is necessary for neuroproteasome-inhibition induced Tau aggregation. Taken together with the dramatic increase in newly synthesized Tau protein following neuroproteasome inhibition, we propose two models to explain our observations (**Fig S7D**): one, that neuroproteasomes degrade newly synthesized Tau, or two, that neuroproteasomes indirectly regulate the translation machinery itself. The latter is unlikely because only ∼500 proteins increase after neuroproteasome inhibition, which suggests some degree of specificity of what is regulated by neuroproteasomes. Based on the small number of neuroproteasome substrates (*4*) and even smaller number of proteins which we identify in this study which become insoluble, we suggest that the model that neuroproteasomes degrade Tau to be the most likely option. While further experiments are needed to determine which is accurate, our observation that newly synthesized and endogenous Tau forms aggregates following neuroproteasome inhibition is likely to be an important finding for the field. Indeed, if newly synthesized Tau is a neuroproteasome substrate, future analysis would determine if Tau mutations or Tau seeding would perturb the balance of protein synthesis and degradation. This would be an important cautionary consideration for using these common models to study neuroproteasome-dependent Tau aggregation.

As a concluding note, our findings increase the urgency to understand fundamental aspects of neuroproteasome biology. For example, revealing the genetic and cell biological elements which control neuroproteasome localization and trafficking are likely to be critical to reveal how ApoE exerts an effect on neuroproteasome localization. The majority of the other components identified are involved in synaptic vesicle recycling and membrane trafficking (NSF, COPI, Vps35 and Sorl1, AP2 and AP3). We believe this reflects the underlying cellular mechanisms for regulating localization of these complexes to the plasma membrane, a subject for future study. Understanding these, as well as related mechanisms like how newly synthesized proteins are degraded by neuroproteasomes, are likely to open up therapeutic avenues that can impinge on neuroproteasome biology to boost neuroproteasome dependent-proteostasis and delay or prevent protein aggregation.

## Supporting information

Supplemental Figures Compiled

## Acknowledgements

We thank the families and patients who contributed human tissue which enabled the discoveries in this manuscript. We thank Peter Davies for the DA9 antibody and Virginia Lee for the R2295 antibody. We thank Abid Hussaini for providing the hTau-KI mice and Carol Troy for the perfusion setup. We thank Claire Chen for IHC training, Ottavio Arancio and Elentina Argyrousi for stereotactic surgery training, and Rejji Kuruvilla and Guillermo Moya Alvarado for technical assistance with the FLAG antibody feeding experiments. We also thank Leona Lee for help with plasmid sequence alignments. We thank Ulrich Hengst and Clarissa Waites for discussions, Leah Cairns, Nikhil Sharma, Franck Polleux, and Lloyd Greene for careful reading and comments, and Sadie for invaluable laboratory support. We also thank Michael Shelanski, Richard Mayeux, and the Taub Institute for support and start-up funding. We thank the Thermo Fisher Scientific Center for Multiplexed Proteomics at Harvard Medical School (http://tcmp.hms.edu).

## Author Contributions

KVR conceptualized the project and KVR and VP designed the methodology for all experiments. VP performed the majority of experiments. In addition, MS performed all stereotactic injections and in vivo surgeries, JB performed the proteomic screen in mice, NAS and CN designed and synthesized neuroproteasome-specific inhibitors, RM performed the majority of the IHC sectioning and produced primary embryonic cultures, KDKV performed all Puro-PLA experiments and analysis, XW performed proteasome isolations, JF analyzed proteomic datasets, BTC performed blinded ThioS quantification, JN helped arrange figures, GM performed perfusion experiments for ApoE-TR experiments and blinded experimenters and trained MS in stereotactic surgeries, MSt prepared lipidated ApoE particles, HF maintained mouse crosses, DA, JN, XW, RM, CN, MS, and VP aided with writing and KVR did the majority of writing. BH provided ApoE human postmortem samples and important insights, DMH provided reagents for the in vitro lipidated ApoE experiments and important insights, and TN provided ApoE2, 3, and 4-KI mice as well as hTau/ApoE2, 3, and 4 double KI mice. KVR was responsible for acquiring funding and supervised the project.

