## Supplemental Figures Compiled for "Dysregulation of neuroproteasomes by ApoE isoforms drives endogenous Tau aggregation"

### Supplementary Figures

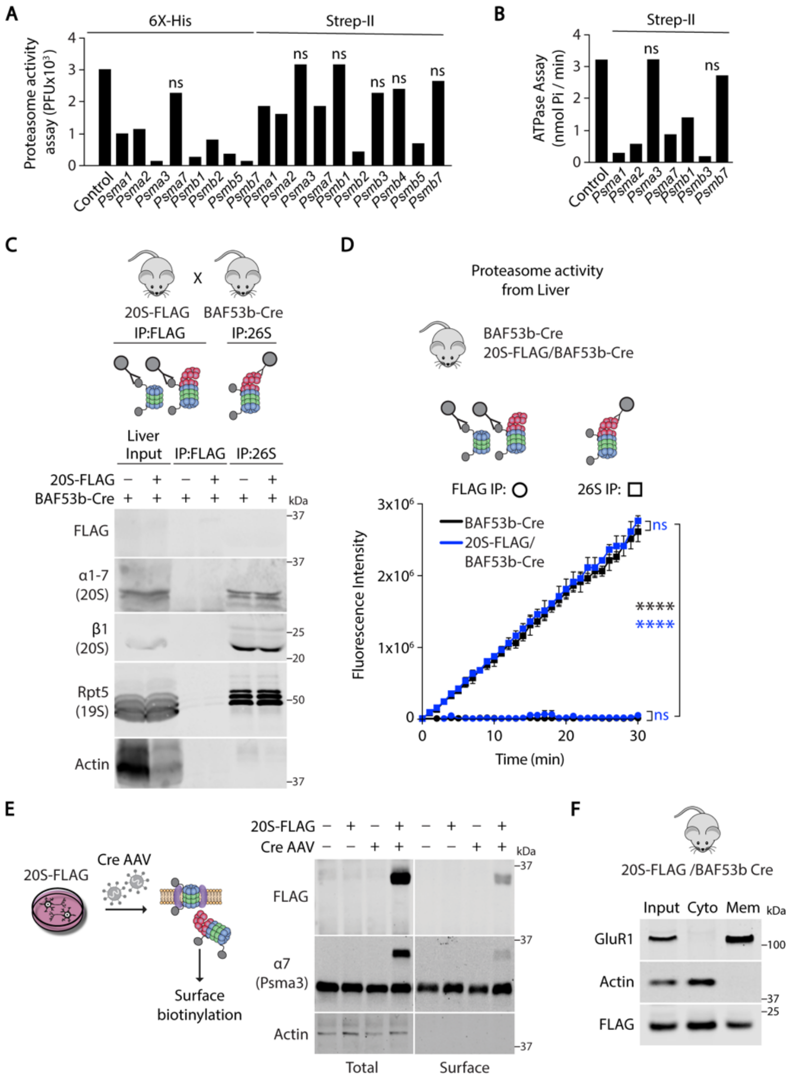


Figure S1
Paradise et al, 2023

**Supplemental Figure 1: Neuroproteasomes co-purify with ApoE and Lrp1**

**(A)** Assessment of catalytic activity of epitope-tagged proteasomes in HEK293T cells. Cells were transfected with candidate constructs with epitope tags on the C-terminus of 11 20S subunits, and the 20S proteasomes were purified using Enzo Proteasome Purification Matrix. Proteasome catalytic activity was assessed by monitoring degradation of Suc-LLVY-AMC. Proteasome-dependent cleavage of the substrate increases AMC fluorescence. *$p>0.5$ indicates not significant from control, one-way ANOVA

**(B)** Candidate hits from (a) subjected to ATPase assay to measure the function of the 19S. *$p>0.5$ indicates not significant from control.

**(C)** Proteasome purifications from the cytosol of livers from 20S-FLAG/BAF53b-Cre mice. (Top) 20S-FLAG mice crossed with pan-neuronal Cre driver BAF53b-Cre mice to induce 20S-FLAG expression in neurons. (Middle) Diagram of proteasomal complexes isolated by different affinity methods; Immunoprecipitation (IP) using FLAG beads isolate FLAG-tagged 20S core particle as well as the 20S-containing 26S particle, whereas IP against the 19S only isolates 26S particles. (Bottom) FLAG and 26S IP from mouse liver cytosolic fractions immunoblotted using indicated antibodies. Note, compared to 1(b) which was done from brain cytosol.

**(D)** Catalytic activity of proteasomes isolated by 26S IP (squares) and FLAG IP (circles) from whole liver cytosol. Proteasomes were isolated using FLAG IP from BAF53b-Cre mice (black) or 20S-FLAG/BAF53b-Cre mice (blue). Catalytic activity was assessed by monitoring cleavage of model-substrate, Suc-LLVY-AMC. Data are mean ± SEM from two technical replicated. ****p<0.0001 by Two-Way ANOVA Tukey's Multiple Comparison Test.

**(E)** Validation of inducible 20S-FLAG transgene in primary neurons. (Top) Primary neurons obtained from 20S-FLAG mice were transduced with Cre AAVs at DIV2 to drive 20S-FLAG transgene expression and were subjected to surface biotinylation at DIV14. (Bottom) Lysates (Total) and Streptavidin pulldowns (Surface) were immunoblotted using the indicated antibodies.

**(F)** Validation of membrane fractionation protocol. Whole brains from 20S-FLAG/BAF53b-Cre mice were subjected to membrane fractionation and the lysate (Input), cytosolic fraction (Cyto), and membrane fraction (Mem), were immunoblotted using antibodies against FLAG, a cytosolic protein (Actin), and a surface protein (GluR1).

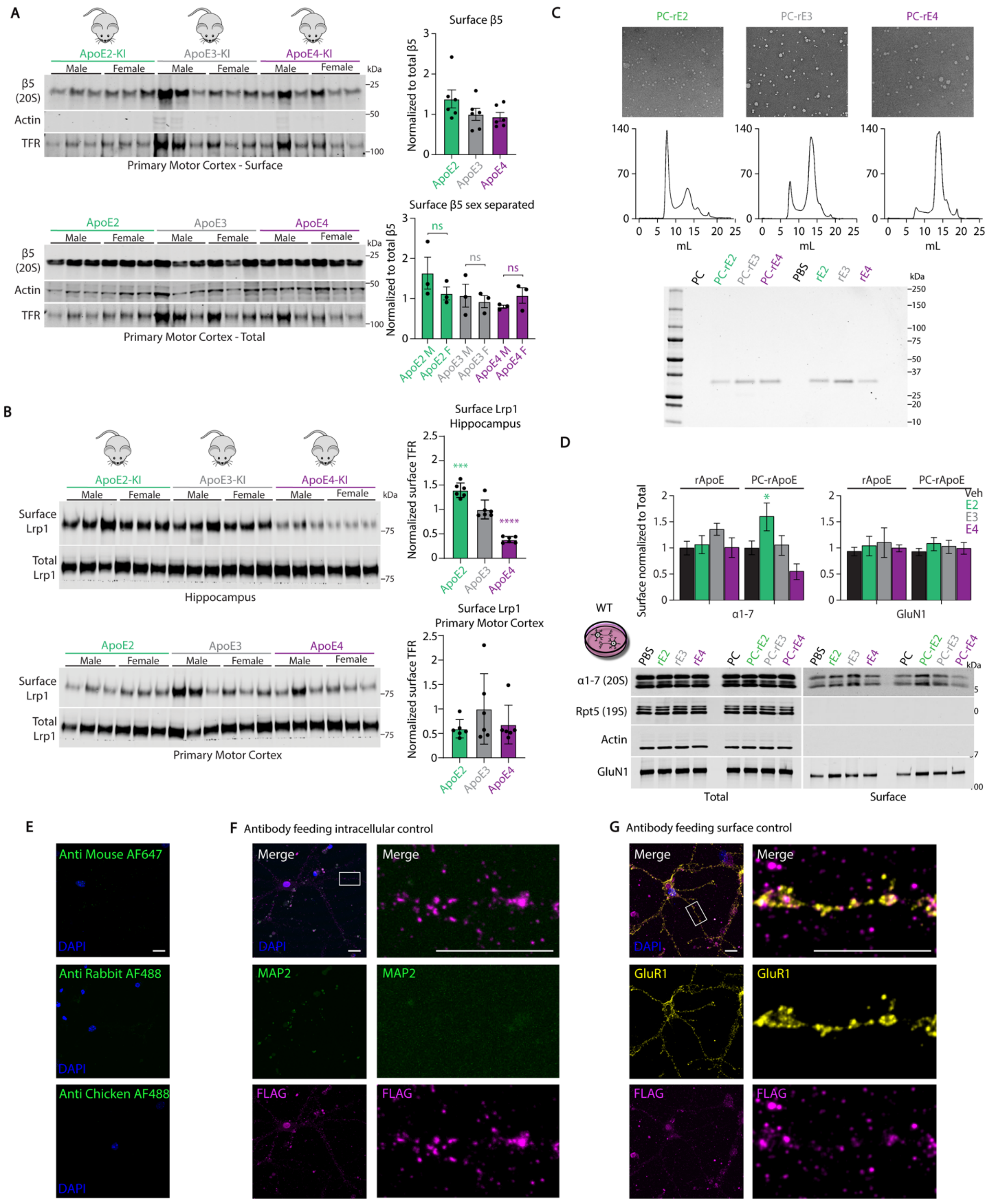


Figure S2
Paradise et al, 2023

**Supplemental Figure 2: ApoE isoforms differentially modulate neuroproteasome localization in primary neurons and *in vivo***

**(A)** Surface biotinylation of the primary motor cortex from hApoe-KI mice to assess neuroproteasome localization. Primary motor cortex (PMC) from ApoE2-KI (green), ApoE3-KI (gray), and ApoE4-KI (purple), male (M) and female (F) mice were subjected to surface biotinylation and streptavidin pulldown to isolate surface exposed proteins. Lysate (Total) and streptavidin pulldowns (Surface)were immunoblotted using indicated antibodies. Quantification of surface β5 intensity was normalized to total β5 intensity. Data shown normalized to ApoE3-KI, N=6 biological replicates per genotype (3 M, 3 F), $p>0.05$ by One-Way ANOVA, Two-Way ANOVA Tukey's Multiple Comparisons Test. Experimenters were blinded to genotype.

**(B)** Representative images from negative stain EM (top) of hApoE lipoproteins PC-rE2 (green), PC-rE3 (gray), and PC-rE4 (purple) and their corresponding chromatograms from size-exclusion chromatography (bottom). (Right) Coomassie G250 stained gel comparing PC-ApoE isoforms and non-lipidated recombinant ApoE isoforms.

**(C)** Antibody feeding of WT neurons treated with exogenous ApoE isoforms. DIV2 primary cortical neurons from 20S-FLAG mice were transduced with Cre AAVs to drive 20S-FLAG transgene expression. At DIV13, neurons were treated with exogenous unconjugated recombinant ApoE isoforms (400 nM; rE2: green; rE3: gray; rE4: purple), or POPC/Cholesterol (PC)-conjugated recombinant ApoE isoforms (400nM; PC-rE2: green; PC-rE3: gray; PC-rE4: purple) for 24 hours and subjected to surface biotinylation. Lysates (Total) and streptavidin pulldowns (Surface) were immunoblotted using indicated antibodies. Quantification of surface α1-7 and GluN1 intensities normalized to corresponding total signal. Data (right) are mean ± SEM normalized to corresponding vehicle controls. N=3 biological replicates, *$p<0.05$, by One-Way ANOVA Tukey's Multiple Comparisons Test.

**(D)** Secondary-only controls for 20S-FLAG antibody feeding experiments. DIV14 primary hippocampal neurons obtained from 20S-FLAG mice received the same treatment as antibody feeding samples as in 2(d), but only secondary antibodies were used. Scale bars=10 μm.

**(E)** Surface protein control for 20S-FLAG antibody feeding experiments. FLAG and GluR1 (surface control) antibodies were fed onto live DIV14 primary hippocampal neurons obtained from 20S-FLAG mice. Scale bars=10 μm.

**(F)** Antibody permeability control for 20S-FLAG antibody feeding experiments. FLAG and MAP2 (intracellular control) antibodies were fed onto live DIV14 primary hippocampal neurons obtained from 20S-FLAG mice. Scale bars=10 μm.

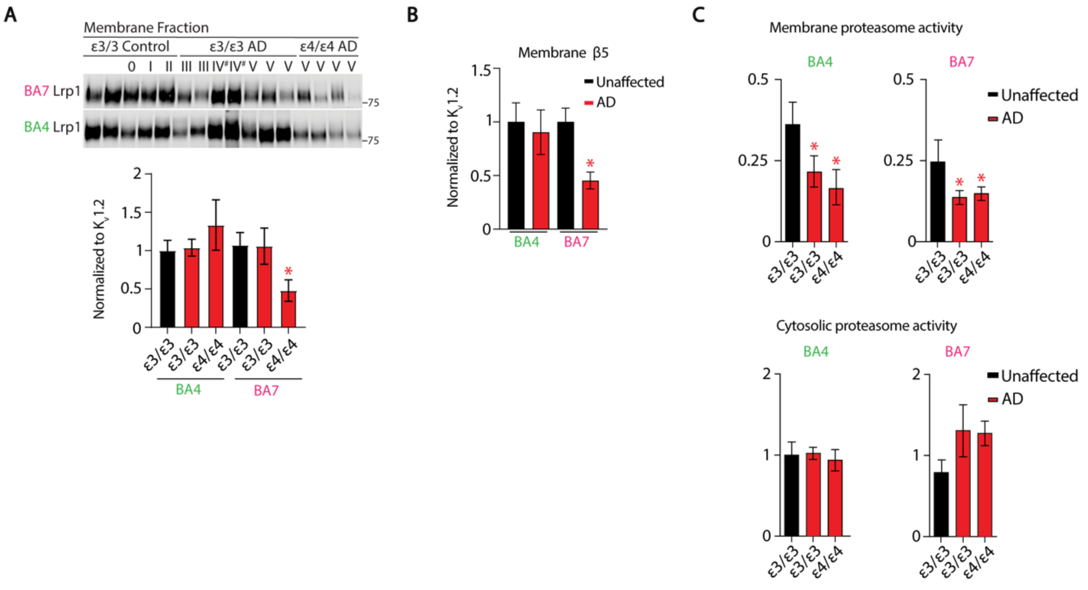


Figure S3
Paradise et al, 2023

**Supplemental Figure 3: Neuroproteasome localization is reduced in AD-vulnerable brain regions in the human brain and is further reduced by ApoE4 genotype**

**(A)** Determining effect of AD and ApoE genotype on Lrp1 membrane localization in BA7 (pink). Membrane fractions from BA7 (pink) and BA4 (green) from Apoε3/3 patients with AD (AD) and without AD (unaffected) and Apoε4/4 patients with AD were immunoblotted using indicated antibodies. Braak stages indicated above. (Bottom) Quantification of Lrp1 intensities were quantified and normalized to membrane loading control $K_V1.2$. Data are mean ± SEM normalized to ε3/3 unaffected. # indicates samples excluded from analysis on basis of high abnormal actin and $K_V1.2$ signal (Fig 3D). **p<0.01 by Two-Way ANOVA Fisher's LSD Test relative to BA7 ε3/3 unaffected.

**(B)** Data from Fig 3D and 3E, analyzed by isolating effect of AD on neuroproteasome membrane localization in BA4 (green) and BA7 (pink). Note that BA7 is more susceptible to AD pathology than BA4.

**(C)** Proteasome catalytic activity of membrane and cytosolic fractions from brain tissues of AD patients of ε3/3 and ε4/4 genotypes in BA4 and BA7. Proteasome activity was assessed by monitoring degradation of Suc-LLVY-AMC. Proteasome-dependent cleavage of the substrate increases AMC fluorescence. Data (right) are mean ± SEM normalized to cytosolic ε3/3 unaffected. *p<.05 by Two-Way ANOVA, Fisher's LSD Test.

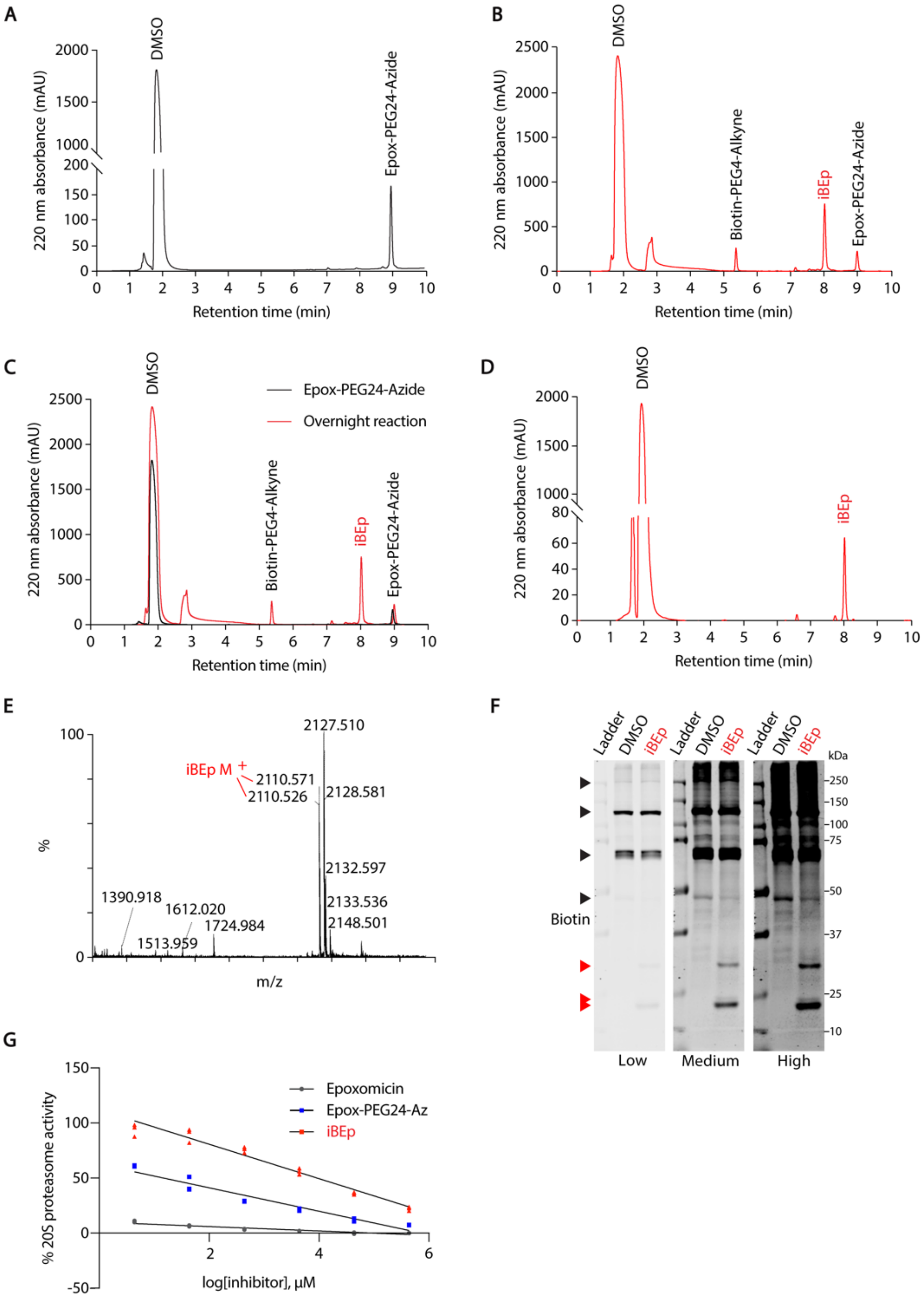


Figure S4
Paradise et al, 2023

**Supplemental Figure 4: Quantitative proteomics reveals that selective inhibition of neuroproteasomes induces accumulation of sarkosyl-insoluble Tau**

**(A)** HPLC analysis for Epox-PEG24-azide.

**(B)** HPLC analysis for the overnight reaction between Biotin-PEG4-alkyne and Epox-PEG24-azide.

**(C)** Overlay of S4A (black) and S4B (red). Note the emergence of peak at 8 minutes corresponding to the formation of iBEp (red) after overnight reaction.

**(D)** HPLC analysis of iBEp after purification.

**(E)** LC/MS spectrum of purified iBEp. Expected mass of iBEp (2110.54) is observed. Other peaks correspond to adducts resulting from LC/MS ionization in the spectrometer.

**(F)** Three exposures (Low, Medium, High) of same blot from primary neurons treated with DMSO or iBEp for 12 hours. Black arrowheads denote endogenous biotinylated proteins, Red arrowheads denote covalent modification of proteasome subunits with biotinylated epoxomicin (iBEp). Note there are All subunits run expected at MW based.

**(G)** Proteasome catalytic activity from HEK293T lysate comparing inhibitory effects of a dose curve of iBEp (red), iBEp precursor Epox-PEG24-Az (blue), and Epoxomicin (dark gray). Catalytic activity was assessed by monitoring degradation of Suc-LLVY-AMC; proteasome-dependent cleavage of the substrate increases AMC fluorescence. N=3 replicates.

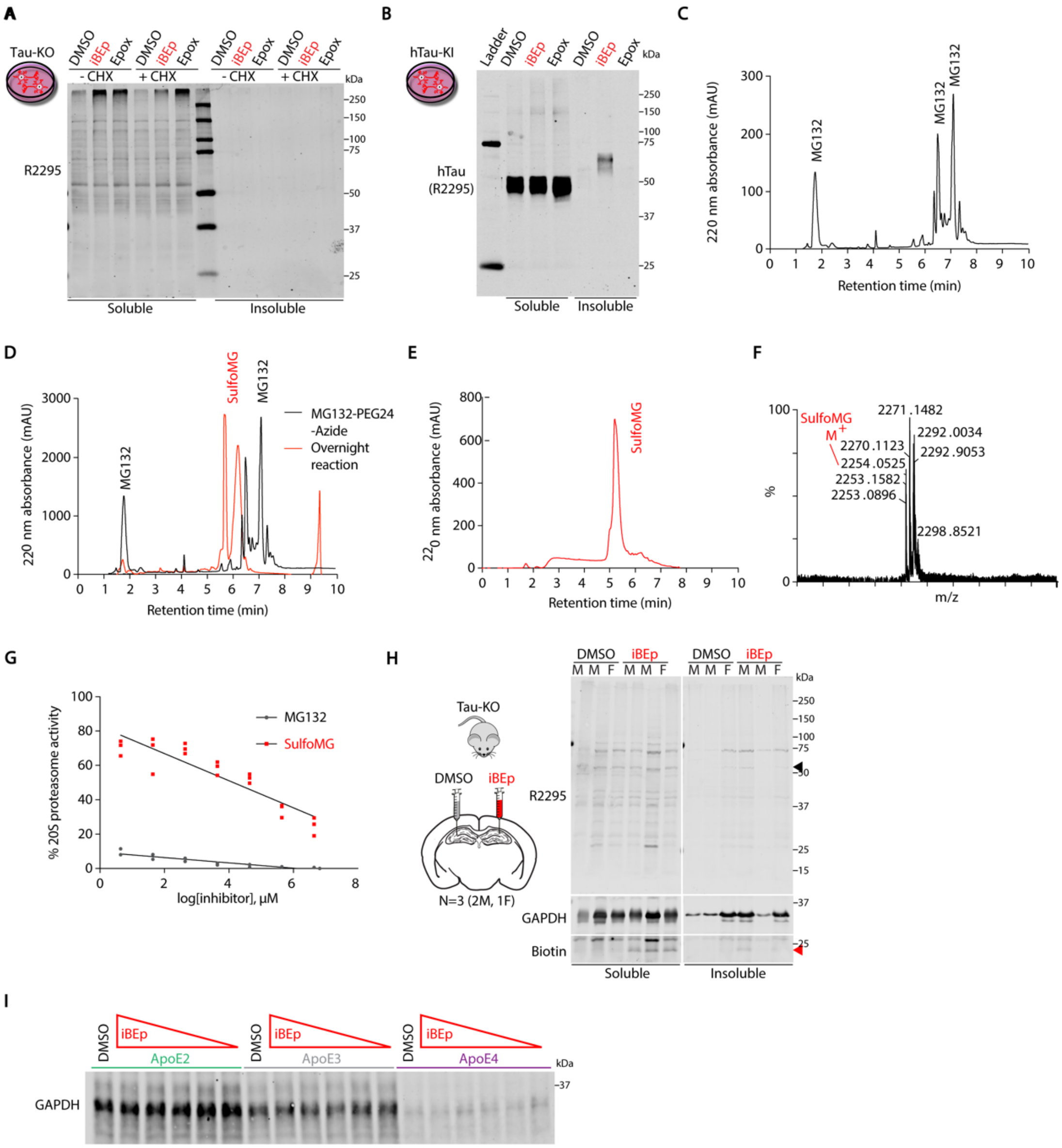


Figure S5
Paradise et al, 2023

**Supplemental Figure 5: Neuroproteasomes are differentially regulated by ApoE isoforms to determine formation of endogenous sarkosyl-insoluble Tau inclusions**

**(A)** Validation of anti-Tau antibody R2295 specificity in primary neurons from Tau knockout (Tau-KO) mice. DIV14 Tau-KO primary cortical neurons were treated as described in (d) with iBEp or Epoxomicin (Epox) and were subjected to sarkosyl fractionation. Sarkosyl-soluble (Soluble) and Sarkosyl-insoluble (Insoluble) fractions were immunoblotted using anti-total Tau antibody R2295. We do not observe any signal at the expected molecular weight for Tau as indicated by the black arrow.

**(B)** Full-length blot from Figure 5B do demonstrate molecular weight of sarkosyl-insoluble Tau species formed following neuroproteasome inhibition with iBEp (red), compared to Epoxomicin (Epox) or DMSO controls.

**(C)** HPLC analysis of MG132-PEG24-Azide.

**(D)** Overlay of HPLC chromatograms of MG132-PEG24-Azide (black) and the overnight reaction between MG132-PEG24-Azide and DBCO-Sulfo-Link-Biotin (red). Note peaks emerging at 5.5 and 6 minutes corresponding to the formation of Sulfo-MG132 (SulfoMG).

**(E)** HPLC analysis of SulfoMG after purification.

**(F)** LC/MS spectrum of purified SulfoMG. Expected mass of SulfoMG (2254.0525) is observed. Other peaks correspond to adducts resulting from LC/MS ionization in the spectrometer.

**(G)** Proteasome catalytic activity from LentiX293T lysate comparing inhibitory effects of a dose curve of SulfoMG (red) and MG132 (dark gray). Catalytic activity was assessed by monitoring degradation of Suc-LLVY-AMC; proteasome-dependent cleavage of the substrate increases AMC fluorescence.

**(H)** Tau-KO Mice (N=3, 2M, 1F) were injected with iBEp (red) in the CA1 region of the hippocampus of the left hemisphere and DMSO was injected contralaterally. The hippocampi were subjected to sarkosyl extraction 72 hours post injection. The sarkosyl-soluble (Soluble) and sarkosyl-insoluble (Insoluble) fractions were immunoblotted using indicated antibodies and fluorescent Streptavidin. We do not observe any signal at the expected molecular weight for Tau as indicated by the black arrow. Red arrow indicates iBEp signal.

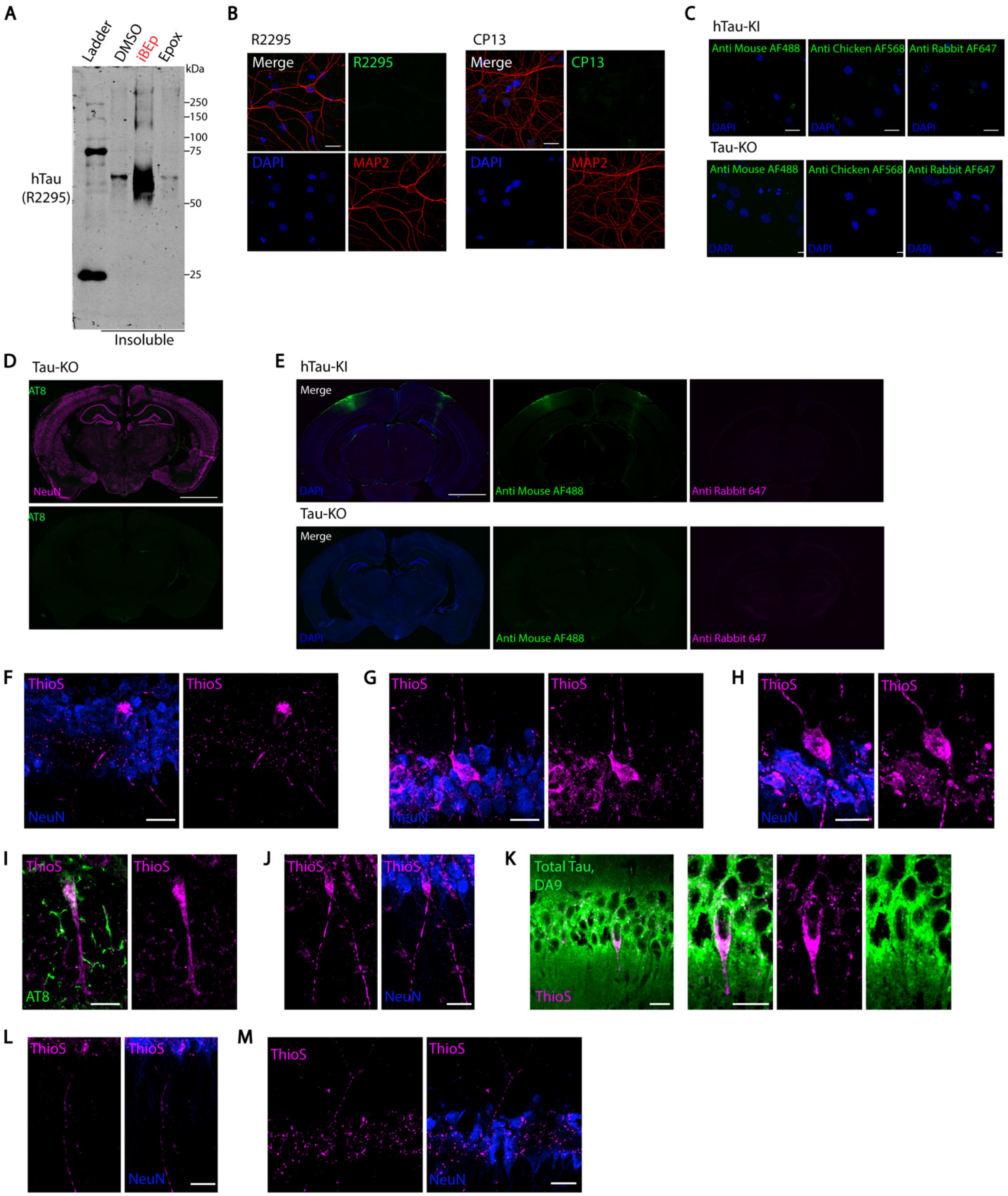


Figure S6
Paradise et al, 2023

**Supplemental Figure 6: Neuroproteasome inhibition induces endogenous phosphorylated, sarkosyl-insoluble, Thioflavin S-positive Tau aggregates**

**(A)** Full-length blot demonstrating high molecular species of sarkosyl-insoluble Tau species formed following neuroproteasome inhibition with iBEp (red), compared to Epoxomicin (Epox) or DMSO controls.

**(B)** Immunocytochemical validation of specificity of R2295, CP13, and AT8 antibodies. DIV14 primary hippocampal neurons obtained from Tau knockout (Tau-KO) mice were stained using indicated antibodies and DAPI. Scale bars=20μm.

**(C)** Secondary only controls for immunocytochemical analysis of CP13 accumulation. DIV 14 primary hippocampal neurons obtained from hTau-KI (top) and Tau-KO (bottom) mice were stained using only secondary antibodies as indicated. Scale bars=20μm.

**(D)** Immunohistochemical validation of specificity of AT8 antibody. Sections obtained from Tau-KO mice were stained using indicated antibodies. Scale bars=1mm.

**(E)** Secondary only controls for immunohistochemical analysis of AT8 accumulation. Sections obtained from hTau-KI mice, injected with iBEp in the CA1 region of the hippocampus of the left hemisphere and DMSO contralaterally (top) and Tau-KO mice (bottom) were stained using indicated secondary antibodies and DAPI. Scale bars=1mm.

**(F-H)** Immunohistochemical analysis of mice stereotactically injected with iBEp to measure Thioflavin-S positive Tau aggregates *in vivo*. Four-to-five month old hTau-KI mice were treated identically to (c), but stained for Thioflavin S (magenta) to visualize β-sheet containing aggregates and immunostained with NeuN (blue). N=7 (3M, 4F). Each panel is separate representative examples of flame-like Thioflavin-S positive inclusion in iBEp-treated hippocampi. Scale bar=20μm.

**(I)** Hippocampal sections from iBEp-injected mice co-stained with Thioflavin S (magenta) and AT8 (green), overlap appears white. Contains representative example of flame-like ThioS+ inclusions with additional thread-like ThioS staining. Images are single Z-plane images to accurately demonstrate co-localization of ThioS and AT8 staining. Scale bar=20μm.

**(J)** Hippocampal sections from iBEp-injected mice co-stained with Thioflavin S (magenta) and NeuN (blue). Scale bar=20μm. Contains representative example of neuron in CA1 with flame-like ThioS+ inclusion with additional thread-like ThioS staining in hippocampal neuropil. Scale bar=20μm.

**(K)** Hippocampal sections from iBEp-injected mice co-stained with Thioflavin S (magenta) and total Tau DA9 (green). Scale bar=20μm. Contains representative example of neuron in CA1 with flame-like ThioS+ inclusion. Images are single Z-plane images to accurately demonstrate co-localization of ThioS and AT8 staining. Scale bar=20μm.

**(L, M)** Hippocampal sections from iBEp-injected mice co-stained with Thioflavin S (magenta) and NeuN (blue). Scale bar=20µm. Contains representative example delicate thread-like ThioS staining in hippocampal CA1 neuropil. Scale bar=20µm.

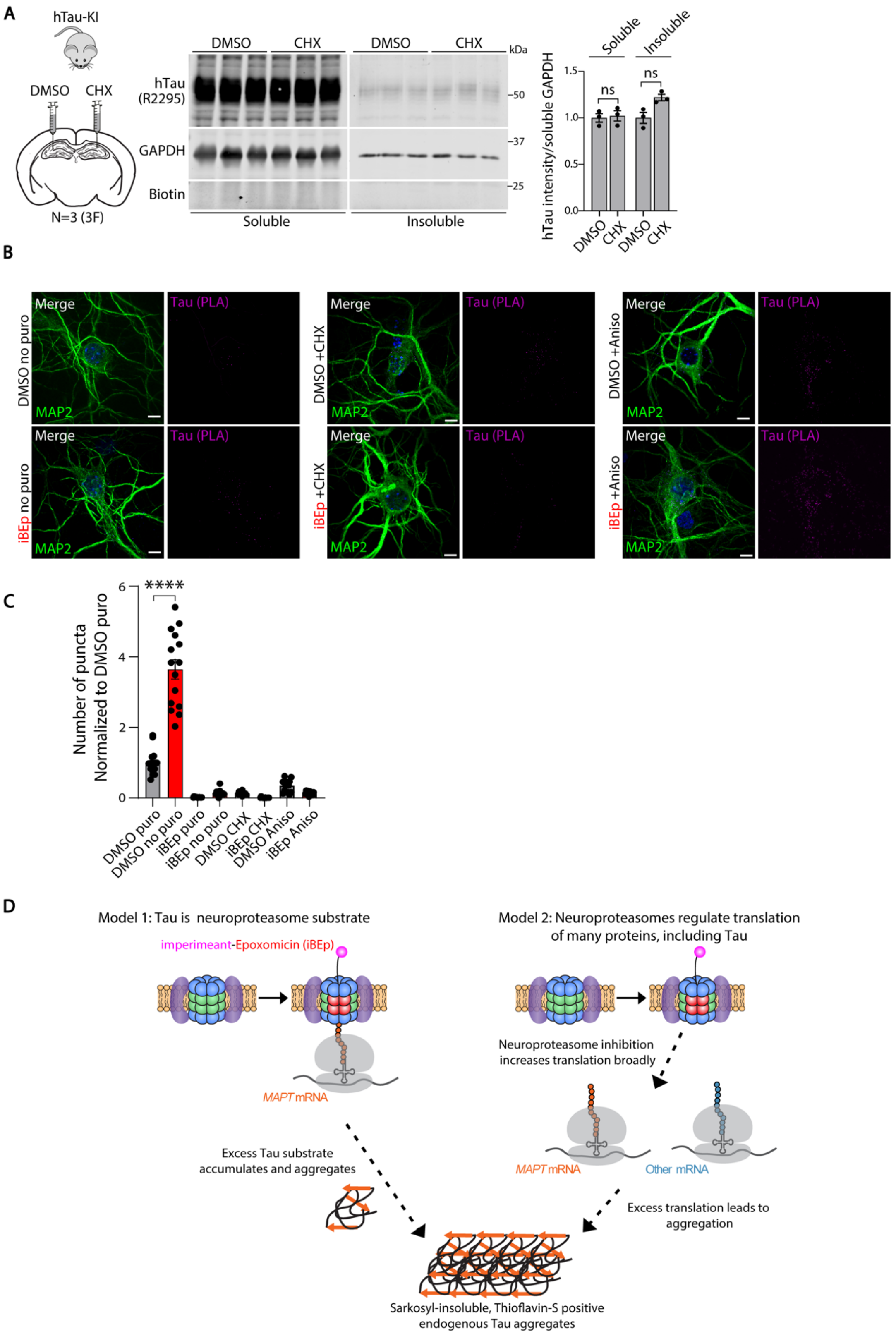


Figure S7
Paradise et al, 2023

**Supplemental Figure 7.** ***De novo*** **protein synthesis is required for neuroproteasome-dependent induction of endogenous Tau aggregates**

**(A)** Four-to-five month old hTau-KI mice were stereotactically injected with CHX in the CA1 region of the hippocampus of the left hemisphere and 0.07% DMSO was injected contralaterally. The hippocampi were subjected to sarkosyl fractionation 72 hours post injection. The sarkosyl-soluble (Soluble) and sarkosyl-insoluble (Insoluble) fractions were immunoblotted using indicated antibodies and fluorescent streptavidin. Quantification of soluble and insoluble Tau intensities were normalized to soluble GAPDH. Data shown (right) are mean ± SEM normalized to respective DMSO controls. N=3 biological replicates (3F), ns indicates not significant by Paired T-test.

**(B)** Primary DIV14 hippocampal neurons treated with DMSO or iBEp (red) were either 1) not puromycylated (No Puro), 2) puromycylated for 10 minutes in the presence of cycloheximide (CHX), or 3) puromycylated for 10 minutes in the presence of anisomycin (Aniso). Tau-PLA-Puro labeling (pink) denotes newly synthesized Tau puncta, total Map2 positive dendrites (green), DAPI (blue).

**(C)** Quantification of Tau-PLA-Puro signal intensity from (B) and from Figure 7H were plotted together and normalized to DMSO Puro. ****p<0.0001 by Paired T-test. Scale bar= 5µm.

**(D)** Model schematic
